# Microglial ferroptotic stress causes non-cell autonomous neuronal death

**DOI:** 10.1101/2022.04.28.489869

**Authors:** Jeffrey R. Liddell, James B.W. Hilton, Kai Kysenius, Sara Nikseresht, Lachlan E. McInnes, Dominic J. Hare, Bence Paul, Benjamin G. Trist, Kay L. Double, Stephen W. Mercer, Scott Ayton, Blaine R. Roberts, Joseph S. Beckman, Catriona A. McLean, Anthony R. White, Paul S. Donnelly, Ashley I. Bush, Peter J. Crouch

## Abstract

**Background:** Ferroptosis is a form of regulated cell death characterised by lipid peroxidation as the terminal endpoint and a requirement for iron. Although it protects against cancer and infection, ferroptosis is also implicated in causing neuronal death in degenerative diseases of the central nervous system (CNS). The precise role for ferroptosis in causing neuronal death is yet to be fully resolved.

**Methods:** To elucidate the role of ferroptosis in neuronal death we utilised co-culture and conditioned medium transfer experiments involving microglia, astrocytes and neurones. We ratified clinical significance of our cell culture findings via assessment of human CNS tissue from cases of the fatal, paralysing neurodegenerative condition of amyotrophic lateral sclerosis (ALS). Finally, we utilised the SOD1^G37R^ mouse model of ALS and a novel CNS-permeant ferroptosis inhibitor to verify pharmacological significance *in vivo*.

**Results:** We found that sublethal ferroptotic stress selectively affecting microglia triggers an inflammatory cascade that results in non-cell autonomous neuronal death. Central to this cascade is the conversion of astrocytes to a neurotoxic state. We show that spinal cord tissue from cases of ALS exhibits a signature of ferroptosis that encompasses atomic, molecular and biochemical features. Moreover, a molecular correlation between ferroptosis and neurotoxic astrocytes evident in ALS-affected spinal cord is recapitulated in the SOD1^G37R^ mouse model where treatment with the novel, CNS-permeant ferroptosis inhibitor, Cu^II^(atsm), ameliorated these markers and was neuroprotective.

**Conclusions:** By showing that microglia responding to sublethal ferroptotic stress culminates in non-cell autonomous neuronal death, our results implicate microglial ferroptotic stress as a rectifiable cause of neuronal death in neurodegenerative disease. As ferroptosis is currently primarily regarded as an intrinsic cell death phenomenon, these results introduce an entirely new pathophysiological role for ferroptosis in disease.

## Background

Disease-modifying treatments for neurodegenerative conditions of the central nervous system (CNS) have proved elusive because the major pathways to neuronal death are yet to be elucidated. Defining and targeting histopathologically conspicuous neuronal features has been a focus in the field. However, non-cell autonomous pathways to neuronal death are also implicated, whereby the glial cells that are essential for healthy neuronal function (1, 2) actively contribute to neuronal death in disease (3, 4). Glial cells, therefore, may represent a more suitable target than neurones for disease-modifying intervention.

The contribution of glial cells to neuronal death is clearly illustrated in amyotrophic lateral sclerosis (ALS), an aggressive adult-onset neurodegenerative disease that results in paralysis and death due to the loss of motor neurones in the brain and spinal cord (5). Restricting expression of ALS-causing mutations specifically to astrocytes results in neuronal death *in vitro* (6-8) and *in vivo* (9), and astrocytes derived from patients with sporadic ALS (i.e. with no known causal mutations) are neurotoxic (10).

*In vitro* studies show that inflammatory activation of microglia – resident immune cells of the CNS – can provoke naturally neurotrophic astrocytes to convert to a neurotoxic state (11). This model system indicates an inflammatory signalling pathway in which the interplay between microglia and astrocytes precedes neuronal death. Parallel evidence from ALS, Parkinson’s, Alzheimer’s, progressive multiple sclerosis and Huntington’s disease indicates that microglia and astrocytes conspiring to cause neuronal death may be a common feature of neurodegeneration (11-13).

Here, we examined whether the iron-dependent form of regulated cell death described as ferroptosis (14) contributes to glial-mediated neuronal death. Although ferroptosis can propagate intercellularly (15), it remains regarded as the terminal endpoint for each cell. Results presented herein provide the first evidence for sublethal ferroptotic stress initiating an inflammatory cascade that culminates in non-cell autonomous neuronal death. Furthermore, although ferroptosis is implicated in neurodegenerative disease (16, 17), its pathophysiological role is currently unproven. Our results from human and mouse spinal cord tissue indicate pertinence of glial ferroptotic stress as a cause of neuronal death in neurodegenerative disease.

## Materials and Methods

### Preparation of primary cell cultures

Pregnant C57BL/6JArc females obtained from the Australian Animal Resource Centre were used for generating the neonates used for primary glial cultures and the E14-E15 embryos used for primary neuronal cultures.

#### Primary mixed glial culture

Mixed glial cultures were prepared according to the method of Hamprecht and Loeffler (18) as previously described (19). Newborn pups were decapitated then their brains removed and placed in preparation buffer (137 mM NaCl, 5.35 mM KCl, 0.22 mM KH_2_PO_4_, 0.17 mM Na_2_HPO_4_, 58.5 mM sucrose, 5.55 mM glucose, 200 U/mL penicillin, and 200 µg/mL streptomycin). Brains were sequentially passed through 250 µm and 135 µm nylon gauze, then centrifuged at 500 RCF for 5 min. Dissociated cells were resuspended in DMEM containing 10% foetal bovine serum, 20 U/mL penicillin and 20 µg/mL streptomycin, and plated at 150,000 cells/cm^2^. Cells were maintained at 37°C in 10% CO_2_ for at least 2 weeks, and media was renewed every 7 days.

#### Primary microglia culture

Microglia were harvested from mixed glial cultures according to the method of Saura *et al*. (20), as previously described (19). Briefly, mixed glial cultures were washed in Dulbecco’s modified Eagle’s medium (DMEM) before incubation in a 5:1 mix of DMEM and trypsin-EDTA (Sigma #T4174). After detachment of astrocytes, adherent microglia were either washed in DMEM before medium replaced with mixed glial conditioned medium, or scraped, collected, and re-plated in mixed glial-conditioned media at 20,000 cells/cm^2^. For the later, medium was changed 7-12 h later to Iscove modified Dulbecco media (IMDM) supplemented with 10% FBS and penicillin/streptomycin. Microglial cultures were maintained at 37°C in 5% CO_2_ for 2 days before experiments.

#### Primary neurone culture

Neurones were harvested from the cortices of mouse embryos at E14-E15 based on procedures previously described (21), plated at 250,000 cells/cm^2^ and maintained in neurobasal medium supplemented with 2% B27, 1 mM glutamine, and 0.1% gentamycin at 37°C in 5% CO_2_ for 6 days before experiments. Cytosine arabinoside (2 µM) was added after 1-2 days in culture. For (1*S*,3*R*)-methyl-2-(2-chloroacetyl)-2,3,4,9-tetrahydro-1-[4- (methoxycarbonyl)phenyl]-1*H*-pyrido[3,4-b]indole-3-carboxylic acid (RSL3) challenge experiments, neurones were maintained in medium supplemented with B27 without antioxidants for 6 days and treated in neurobasal medium conditioned by mixed glial cultures for 2 days. Where indicated, mixed cultures of neurones and astrocytes were generated by omitting cytosine arabinoside.

#### Primary astrocyte culture

In preliminary RSL3 titration experiments using mixed glial cultures, microglia were almost completely killed at certain concentrations of RSL3 whereas the astrocyte monolayer appeared to be unaffected. After further investigation, we standardised the following conditions to generate astrocyte cultures with microglia removed. Mixed glial cultures were treated with 300 nM RSL3 overnight, followed by washing and 2 days recovery incubation in serum-containing media, before subsequent treatments as indicated. Effectiveness of microglial removal was confirmed by expression of the microglial molecular markers *Cx3cr1* and *C1qa* by quantitative reverse transcription polymerase chain reaction (RT-PCR), and live-cell imaging. From the latter, the total number of cells per field of view was determined by counting Hoescht-positive nuclei, averaged from 9 fields per condition. The proportion of cells represented by microglia was derived from the morphology of nuclei, isolectin staining and cell mobility. RSL3 toxicity in normal mixed glial cultures was compared to astrocyte cultures with microglia removed (assessed by 3- (4,5-dimethylthiazol-2-yl)-2,5-diphenyltetrazolium bromide (MTT) and lactate dehydrogenase (LDH) assays after 24 h RSL3 treatment) to confirm that exposure to 300 nM RSL3 did not induce a persistent change in the subsequent sensitivity of astrocytes to RSL3 toxicity.

### Cell culture experimental treatments

#### Ferroptosis inducers and inhibitors

For all treatments with ferroptosis inducers and inhibitors, cells were treated for 24 h unless otherwise indicated. Unless otherwise indicated, ferroptosis inducers and inhibitors were applied at the following concentrations: erastin, 10 µM; buthionine sulphoxamine (BSO), 1 mM; ferric ammonium citrate (FAC) as an iron source, 100 µM; liproxstatin-1 (Lip1), 100 nM; ferrostatin-1, 100 nM; deferiprone (DFN), 100 µM.

For viability and lipid peroxidation studies, unless otherwise indicated, RSL3 was applied at 2 µM, Cu^II^(atsm) was applied at 1 µM, and cells were pre-treated with BSO or erastin for 24 h before additional treatments. Where microglial cultures were treated with RSL3, viability and lipid peroxidation were assessed in presence or absence of Lip1 (20 nM) or DFN (200 µM). For live cell imaging, mixed glial cultures were treated with 100 nM RSL3 unless otherwise indicated, and 2 µM Cu^II^(atsm).

Where microglia in mixed glial cultures were assessed in response to BSO, cells were treated with BSO for 4 days. Cells also treated with Lip1 in these experiments were co-treated with BSO and Lip1 followed by a second bolus application of Lip1 after 2 days. A single co-treatment of Lip1 was not effective (data not shown). Treating mixed glial cells with BSO for 24 h did not substantially influence microglia cell numbers (data not shown).

For transcript analyses in response to ferroptosis inducers, microglial cultures were treated for 8 h with RSL3 (25 nM) in presence or absence of Lip1 (20 nM) or Cu^II^(atsm) (200 nM). Mixed glial and astrocyte cultures were treated with either RSL3 (up to 400 nM) or erastin plus iron as FAC (10 µM) in the presence or absence of Lip1, DFN or Cu^II^(atsm) (500 nM). When RSL3 was applied in combination with iron, transferrin-bound iron was used (10 µM transferrin, 20 µM iron). Addition of transferrin-bound iron to RSL3 treatments was confirmed to exacerbate RSL3 toxicity and increase lipid peroxidation (data not shown).

#### Glial conditioned media neurotoxicity experiments

Mixed glial, astrocyte and microglial cultures were treated with RSL3 (mixed glial and astrocyte cultures, 100 nM; microglial cultures, 1 nM), erastin, or lipopolysaccharide (LPS; 1 ng/mL), in presence or absence of Lip1 (200 nM) or Cu^II^(atsm) (200 nM) for 24 h in neurobasal medium (conditioned for 2 days by mixed glial cultures for microglial cultures) before the conditioned media was collected. Protease inhibitor (Roche Complete EDTA-free) was added and the media centrifuged (1000 RCF, 5 min) to remove any cells or debris. The glial-conditioned media was concentrated 20-fold using pre-wetted 30 kDa molecular weight cut off Amicon Ultra-15 centrifugal filters (4000 RCF, 10 min). Retentate was diluted with filtrate until equivalent to 4-fold concentration of neat media (thus maintaining the original concentration of small molecular weight species). Diluting concentrated glial-conditioned media with fresh neurobasal medium (thus diluting any unsequestered small molecular weight species still present in the retentate by 5-fold compared to neat media) did not alter results (data not shown).

Neurones were treated for 24 h with the glial-conditioned media neat or as 4-fold concentrated retentate. When the conditioned media was concentrated more than 4-fold, control retentate (mixed glia treated in the absence of stressors) became neurotoxic (data not shown). Neat conditioned media from RSL3 or erastin-treated glial cells induced mild neurotoxicity (data not shown). Neurones were also treated with conditioned media filtrate to confirm lack of toxicity of unsequestered small molecules including the ferroptosis inducers present in conditioned media. The same concentration of stressors (LPS, RSL3, erastin) was also added directly to control conditioned media and applied to neurones. After treatment, neuronal survival was assessed by MTT assay. The LDH assay could not be used as conditioned media contained a sufficient quantity of LDH activity to interfere with the quantification of neuronal LDH activity.

#### Preparation of transferrin-bound iron

Transferrin-bound iron was prepared by combining acidic FeCl_3_ with nitrilotriacetic acid at a 1:10 ratio. This was combined 1:1.4 with 1.4% NaHCO_3_, and then 1:1 with apo-transferrin at half the molar ratio of FeCl_3_ in PBS. The final stock solution was 125 µM transferrin, 250 µM iron, 2.5 mM nitrilotriacetic acid, 18 mM HCl, 0.43% NaHCO_3_, 50% PBS. All experiments with transferrin-bound iron included an equivalent diluent control, which was prepared as above but with FeCl_3_ and apo-transferrin omitted. For all analyses, results from cells treated with diluent did not differ from those treated with control media.

### Live-cell imaging of lipid peroxidation

Cells were treated with Hoescht-33342 (1 µg/mL) to identify nuclei, Dylight 649-labelled isolectin (Vector labs; 2 µg/mL) to identify microglia, and C11-BODIPY (0.5 µM) for lipid peroxidation for 15 min before addition of treatments and imaging in an Operetta high-content imaging system (PerkinElmer) in a temperature and CO_2_-controlled chamber at the indicated time intervals. Fluorescent dyes were imaged as follows: Hoescht-33342 Ex380±40nm/Em410-480nm; isolectin Ex630±20nm/Em640-680nm; reduced C11-BODIPY Ex570±20nm/Em560-630nm; oxidised C11-BODIPY Ex475±30nm/Em500-550nm. Images were captured with Harmony software (PerkinElmer). Image analysis and reconstruction of image sequences into videos was conducted with ImageJ (1.51s) software. To account for potential differences in accumulation of C11-BODIPY, lipid peroxidation was calculated from the ratio of oxidised to reduced C11-BODIPY with background correction. Time-courses of lipid peroxidation were generated from 4-8 imaged fields per condition per biological replicate.

### Cell culture C11-BODIPY lipid peroxidation assay

Cells seeded in 96 well culture plates were co-treated with C11-BODIPY (5 µM) and the indicated treatments. After 3 h, cells were washed twice with PBS and fluorescence quantified from the cell monolayer at Ex581nm/Em596nm for reduced C11-BODIPY, and Ex490nm/Em517nm for oxidised C11-BODIPY in an EnSpire multimode plate reader (PerkinElmer). These excitation and emission wavelengths were empirically determined in control experiments. Fluorescence was quantified from 4 points per well, with triplicate wells per treatment. Lipid peroxidation was calculated as the ratio of oxidised to reduced C11-BODIPY after correcting for background fluorescence.

### Viability assays

Cell survival was determined by 3-(4,5-dimethylthiazol-2-yl)-2,5-diphenyltetrazolium bromide (MTT) reduction. Briefly, following treatments, MTT (12 mM) was added to cells to a final concentration of 480 µM. Cells were incubated for 1 h before media was removed and cells lysed in DMSO and absorbance measured at 562nm.

Cell death was determined by lactate dehydrogenase (LDH) activity using a Cytotoxicity Detection Kit (LDH) (Roche). Briefly, following treatments, aliquots of media were sampled and combined with a reaction mix. The change in absorbance was measured over time at 490nm and compared to total LDH activity of control wells treated with 1% triton X-100.

### RNA extraction and transcript analyses

All reagents were from Thermo Fisher Scientific unless otherwise indicated and used in accordance with manufacturer’s instructions. RNA was isolated from tissue samples or cultured cells using TRI Reagent (Sigma). Contaminating DNA was degraded by treatment of isolated RNA with DNase (Turbo DNA-free Kit). RNA quantity was determined by nanodrop or Qubit RNA HS Assay Kit. cDNA was synthesised using High Capacity cDNA Reverse Transcription Kit.

For human and mouse tissue samples, 25 ng cDNA was pre-amplified for all genes assessed using Taqman PreAmp Master Mix and pooled Taqman Gene Expression Assays. Pre-amplified cDNA was then diluted 20-fold for subsequent analyses. For cell culture samples, 10 ng of cDNA was used per reaction. All samples were run in triplicate per gene.

Quantitative RT-PCR was performed using Taqman Gene Expression Assays and Taqman Fast Advanced Mastermix on a QuantStudio 6 Flex system (Thermo Fisher Scientific). Relative gene expression was determined via the ΔΔ-ct method normalised to *GAPDH* (human tissue), *Gapdh* (mouse tissue) or *Actb* (cell culture) expression. Normalising genes were chosen based on homology/similarity between the resultant gene expression and the corresponding protein expression determined by immunoblotting. Overall transcript signatures represent the average of gene expression z-scores for all genes in a given gene set for each case, animal or culture.

### Cu^II^(atsm) treatment of ALS model mice

Transgenic mice expressing human SOD1 with the G37R mutation (22) were obtained from the Jackson Laboratories (Stock No: 008342) and a colony maintained by breeding SOD1^G37R^ males with C57BL/6JArc females obtained from the Australian Animal Resource Centre. Non-transgenic littermates were used as control animals for all experiments involving SOD1^G37R^ mice.

SOD1^G37R^ mice were treated with Cu^II^(atsm) commencing when the animals were 140 days old. Cu^II^(atsm) was prepared fresh daily by suspending in standard suspension vehicle (SSV; 0.9% (w/v) NaCl, 0.5% (w/v) Na-carboxymethylcellulose, 0.5% (v/v) benzyl alcohol, 0.4% (v/v) Tween-80) then sonicating. Cu^II^(atsm) was administered at 30 mg kg^-1^ body weight by gavage twice daily 7 days week^-1^ and continued until the animals were killed for analysis or until they reached phenotype end-stage. Control groups involved SOD1^G37R^ mice and non-transgenic littermates gavaged with SSV. Treatment groups were balanced for sex of the animals and treatments balanced across litters as best as possible.

The study involved two separate cohorts of animals. The first cohort included animals that were treated and monitored for phenotype progression, continuing until the animals reached phenotype end-stage. The second cohort included animals that were treated as per the first cohort but killed for tissue collection before they reached phenotype end-stage. These animals were killed at 175-195 days old, with all treatment groups balanced for age.

### Phenotype assessment of ALS model mice

ALS-like phenotype of the SOD1^G37R^ mice was assessed as previously described (23). In brief, motor function of the animals was measured using the rotarod assay which involved the accelerating rod paradigm (4-40 rpm for 180 seconds, performed twice for each day of assessment with only the better performance for each animal used in final data analysis). Survival of the animals was determined as the stage of phenotype progression which necessitated humane killing. Specifically, as soon as the self-righting reflex was lost and persisted for more than 10 seconds.

### Human spinal cord tissue

Frozen sections of lumbar spinal cord were obtained from the Victorian Brain Bank (Australia), the MS Society Tissue Bank (UK), MRC London Neurodegenerative Diseases Brain Bank (King’s College, London, UK), the University of Maryland Brain and Tissue Bank, a biorepository of the NIH NeuroBioBank (Maryland, USA), and the Sydney Brain Bank. Samples were stored at -80°C until processed for analysis.

### Spinal cord tissue processing

Human and mouse spinal cord samples were homogenised in tris(hydroxymethyl)-aminomethane-buffered saline (TBS)-based homogenisation buffer to generate TBS-soluble and -insoluble fractions as previously described (23). Prior to homogenisation, human spinal cord samples were dissected to collected grey matter-enriched material. Mouse spinal cords were homogenised whole. Assessment of ferroxidase activity involved further processing of the TBS-insoluble fraction, involving supplementation with 1% (v/v) triton X-100 then centrifuging (18,000 RCF, 4°C, 5 min) to produce triton X-100 soluble extracts. All fractions were assessed for protein content using the Pierce BCA Protein Assay kit, then normalised to a consistent protein concentration by diluting with the appropriate buffers (i.e. TBS-based homogenisation buffer or the TBS-based homogenisation supplemented with Triton X-100).

### Iron analyses

Iron was quantified in samples using inductively coupled plasma-mass spectrometry (ICP-MS). TBS-soluble and -insoluble fractions of spinal cord generated as described above were assessed for iron concentration using ‘microdroplet’ laser ablation-ICP-MS (LA-ICP-MS) as previously described (24). *In situ* quantitation of iron in human spinal cords was performed using laser ablation-ICP-MS (25) utilising spinal cord embedded in Optimal Cutting Temperature compound and cryo-sectioned at 30 μm in the transverse plane.

### Ferroxidase activity assay

Ceruloplasmin ferroxidase activity in triton X-100 extracts was determined as previously described (26, 27). In brief, for each assay run, fresh solutions of 250 μM human apo-transferrin (Sigma) and 1 mM FeSO_4_ were prepared in N_2_-purged dH_2_O to mitigate ferroxidase-independent oxidation of iron. Reaction mixtures in Hepes-buffered saline (50 mM Hepes, 150 mM NaCl, pH 7.2) contained 50 μM apo-transferrin and sample (or triton X-100 supplemented TBS-based homogenisation buffer as vehicle control), then reactions initiated by adding FeSO_4_ to a final concentration of 100 mM. The formation of holo-transferrin was monitored via change in absorbance at 460nm for 5 min at 25°C.

### Glutathione assay

Glutathione was extracted from tissue with 1% sulfosalicylic acid based on a previously reported procedure (28). Aliquots of lysate were combined with a reaction mix to a final concentration of 200 µM NAPDH, 150 µM 5,5’-dithiobis(2-nitrobenzoic acid), 0.1 U glutathione reductase, and 0.5 mM EDTA in 50 mM NaP_i_ buffer (pH 7.5). The rate of 5-thio-2-nitrobenzoate generation was followed at 405nm. Tissue glutathione was normalised to tissue wet weight.

### Tissue C11-BODIPY lipid peroxidation assay

TBS-insoluble human spinal cord samples were supplemented with 500 nM C11-BODIPY then incubated at ambient temperature for 3 min. Triton X-100 was added to a final concentration of 10% (v/v), samples mixed vigorously, then centrifuged (15,000 RCF, 3 min). Fluorescence in soluble extracts was quantified at Ex581nm/Em596nm for reduced C11-BODIPY, and Ex490nm/Em517nm for oxidised C11-BODIPY in an EnSpire multimode plate reader (PerkinElmer). Lipid peroxidation was calculated as the ratio of oxidised to reduced C11-BODIPY after correcting for background fluorescence. Mouse spinal cord samples were assessed as per human samples except the samples were homogenised in TBS-extraction buffer already supplemented with 500 nM C11-BODIPY.

### SDS-PAGE and immunoblotting

Proteins were assessed by western blot following sodium dodecyl sulphate–polyacrylamide gel electrophoresis (SDS-PAGE) resolution using methods previously described (26). Primary antibodies used were raised to detect: ALOX5 (Sigma, HPA013859); GPX4 (Abcam, ab125066); complement component 3 (Sigma, GW20073F); and GAPDH (Cell Signaling Technology, 2118). Detection utilised horseradish peroxidase conjugated secondary antibodies for anti-rabbit IgG (Cell Signaling Technology, 7074) or anti-chicken IgY (Abcam, ab6877) followed by enhanced chemiluminescence (ECL Advance, GE Healthcare). Abundance of proteins of interest was normalised to the loading control GAPDH and expressed relative to control cases/animals.

### GPX4 activity assay

GPX4 activity in TBS-soluble tissue extracts was determined using phosphatidylcholine hydroperoxide (PC-OOH) as the substrate and RSL3 as a GPX4 selective inhibitor. Procedures used were based on existing protocols (29, 30).

#### PC-OOH preparation

L-α-Phosphatidylcholine (PC; 10 mg) was dissolved in 4 mL 3% (v/v) sodium deoxycholate then diluted to 25 mL using 200 mM borate puffer (pH 9.0). 500,000 U lipoxygenase type I was added, then the solution oxygenised by bubbling through 99% O_2_ for 90 min at 37°C with constant stirring. The PC-OOH solution was then loaded onto a Sep-Pak Vac 35cc C18 cartridge (Waters) which had been activated with 70 mL methanol then pre-equilibrated with 70 mL dH_2_O. After loading the sample, the cartridge was washed with a further 70 mL dH_2_O. An initial 10 mL methanol was added and the flow-through discarded. An additional 10 mL methanol was then added, and the PC-OOH containing eluent collected. Aliquots were prepared then stored at -30°C until used.

#### RSL3 pre-incubations

TBS-soluble tissue extracts containing ∼400 μg protein were incubated on ice for 60 min after adding β-mercaptoethanol to a final concentration of 10 mM. Samples were then divided to two equal aliquots. One was supplemented with 16.7 mM RSL3 and the other with an equivalent volume of the RSL3 vehicle solution (DMSO:ethanol at 1:9). These mixtures were incubated at ambient room temperature for 20 min then kept on ice until used in the GPX4 activity assay.

#### Activity assay

Samples pre-incubated with the GPX4 inhibitor RSL3 or its vehicle control were added to reaction mixture containing 3.3 mM glutathione, 1.1 mM NaN_3_, 110 μM NADPH, 330 U glutathione reductase, and 0.22% (v/v) triton X-100. Basal NADPH consumption was monitored at 340nm for 2 min, followed by addition of PC-OOH, then NADPH consumption monitored at 340nm for a further 10-15 min. GPX4 activity was calculated as the rate of RSL3-sensitive, PC-OOH-dependent NADPH consumption min^-1^ mg^-1^ sample protein.

### Histological iron staining

Paraffin-embedded human spinal cord samples sectioned at 5 mm then de-waxed, were prepared for histological assessment based on a protocol previously described (31). In brief, samples were incubated with 7% (w/v) potassium ferrocyanide in 3% (v/v) hydrochloric acid (1 h, 37°C), followed by 5 min incubation in 3.5 μM 3,3’-diaminobenzidine in 0.0015% (v/v) H_2_O_2_. The reaction was quenched using running H_2_O, then the samples counterstained with neural red. Slides were then washed in H_2_O, dehydrated with increasing ethanol concentration, treated with xylene, then coverslipped. Images were captured using a 3DHISTECH Panoramic SCAN II Scanner.

### Data analyses

All statistical analyses were performed using GraphPad Prism. Statistical outliers were assessed using the ROUT method (32). Data are presented as mean ±S.E.M, violin plots (median, ± 25^th^ and 75^th^ percentiles, truncated at min-max values) or z-scores. Significant differences between groups were determined using two-tailed t-tests, one-way ANOVA where multiple comparisons were corrected using Holm-Sidak’s test, or Mantel-Cox survival test. Significance was determined as P<0.05.

## Results

### Markers of ferroptosis are evident in human, sporadic ALS-afflicted tissue

Iron accumulation increases the risk for ferroptosis. While iron elevation in ALS is implicated by MRI studies (33-35), direct evidence is lacking. We utilised laser ablation-inductively coupled plasma-mass spectrometry (24, 25) to provide the first direct evidence for iron accumulation in ALS within the spinal cord grey matter **(Fig. 1a,b; Supplementary Fig. 1a-k)**. The normal tissue partitioning of iron was changed in ALS spinal cord, with accumulation in the tris-buffered saline (TBS)-insoluble fraction of tissue homogenates **(Supplementary Fig. 1l-n)**. These iron changes were accompanied by decreased ferroxidase activity **(Fig. 1c)**, an activity needed for cellular efflux of iron(36, 37), and altered expression of genes associated with iron handling **(Fig. 1d; Supplementary Fig. 2a)**, indicating perturbed iron homeostasis in ALS **(Fig. 1e)**.

**Figure 1.**
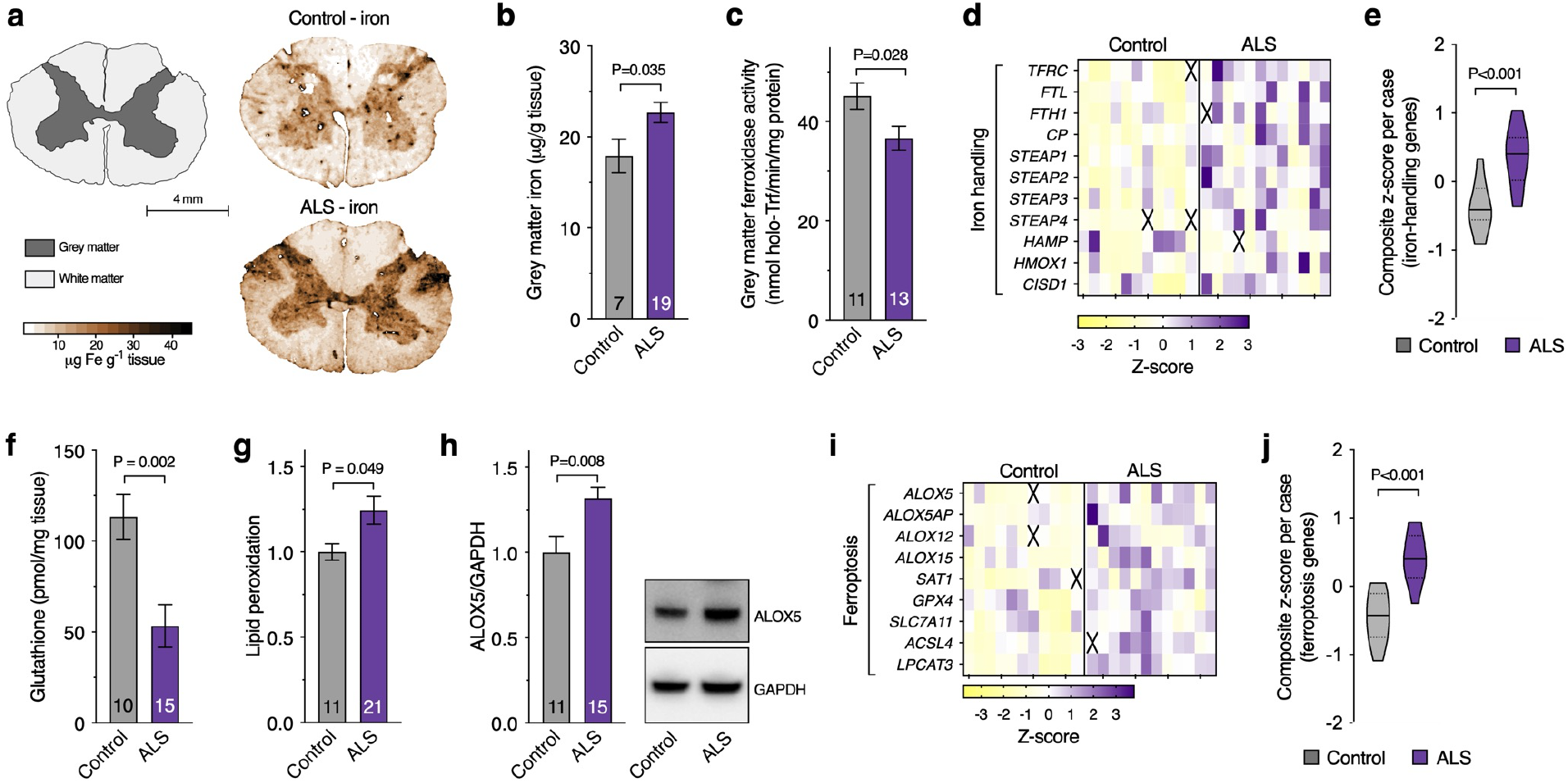
Markers of ferroptosis in human, ALS-affected spinal cord. **(a,b)** Quantitative *in situ* mapping of iron in transverse sections of human spinal cord reveals an overall increase in iron in the ALS-affected grey matter. (**c**) Ferroxidase activity in human, ALS-affected spinal cord. **(d,e**) Transcripts associated with iron handling in human, ALS-affected spinal cord tissue. (**f-h**) Biochemical markers of ferroptosis (glutathione, lipid peroxidation, and ALOX5 protein levels) in human, ALS-affected spinal cord. (**i,j**) Transcripts associated with ferroptosis in human, ALS-affected spinal cord tissue. Data in **g** and **h** are expressed relative to controls. Values in transcript heatmaps (**d,i**) represent z-scores for individual control and ALS cases. Violin plots in **e,j** represent overall transcript signature for features indicated, derived from heatmap data shown in **d,i**, respectively. Crosses in heatmaps represent excluded samples. P values show significant differences where indicated. Numbers in each bar graph represent the number of individual control and ALS cases analysed. Error margins in bar graphs are S.E.M. Solid lines in violin plots represent median, dotted lines represent 25^th^/75^th^ percentiles, truncated at min-max values.

By staunching lipid peroxidation, glutathione peroxidase 4 (GPX4) is a major inhibitor of ferroptosis. The inhibition of GPX4 activity is used to model ferroptosis *in vitro* (38) and its selective ablation from neurones in the murine CNS causes rapid fatality (39). Our analysis of human spinal cord tissue revealed *GPX4* gene expression, GPX4 protein levels, and GPX4 activity were all unchanged in ALS **(Supplementary Fig. 3a-c)**. However, the concentration of glutathione, the substrate for GPX4 ferroptosis checkpoint function, was markedly decreased **(Fig. 1f)**. This was associated with elevated lipid peroxidation in ALS-affected tissue **(Fig. 1g)** and also with elevated levels of ALOX5 protein **(Fig. 1h)**, implicated in the initiation of ferroptosis (40). These findings in tandem with increased aggregated expression of a panel of genes associated with ferroptosis **(Fig. 1i; Supplementary Fig. 2b)** revealed a molecular signature consistent with ferroptosis in ALS **(Fig. 1j)**.

### Ferroptotic stress induces neurotoxic glial activation

Although our quantitation of iron in ALS-affected spinal cord revealed prominent accumulation within the grey matter **(Fig. 1a,b; Supplementary Fig. 1)**, the resolution of this quantitative *in situ* imaging limited identification of cell type specific changes. Perls staining to visualise cellular ferric iron revealed strong staining in non-neuronal cells **(Fig. 2a)**, implying that iron accumulation within ALS grey matter involved accumulation within glia. We therefore examined the potential relationship between glial cells and ferroptosis *in vitro*. Analysis of primary murine cultures of major cell types in the brain revealed that microglia were more vulnerable to the GPx4 inhibitor RSL3 (LD_50_ 1.77 nM) than astrocytes (LD_50_ 496 nM) and neurones (LD_50_ 82.2 nM, **Fig. 2b**). Microglia were also distinctly sensitive to the ferroptosis inducers erastin (inhibitor of the glutamate cystine antiporter) and buthionine sulphoximine (inhibitor of γ-glutamylcysteine ligase required for glutathione synthesis; **Fig. 2c,d**). RSL3 toxicity and lipid peroxidation (detected by the fluorometric lipid peroxidation sensor C11-BODIPY (41)) in microglia were prevented by the ferroptosis inhibitors liproxstatin-1 and deferiprone (**Fig. 2e,f**).

**Figure 2.**
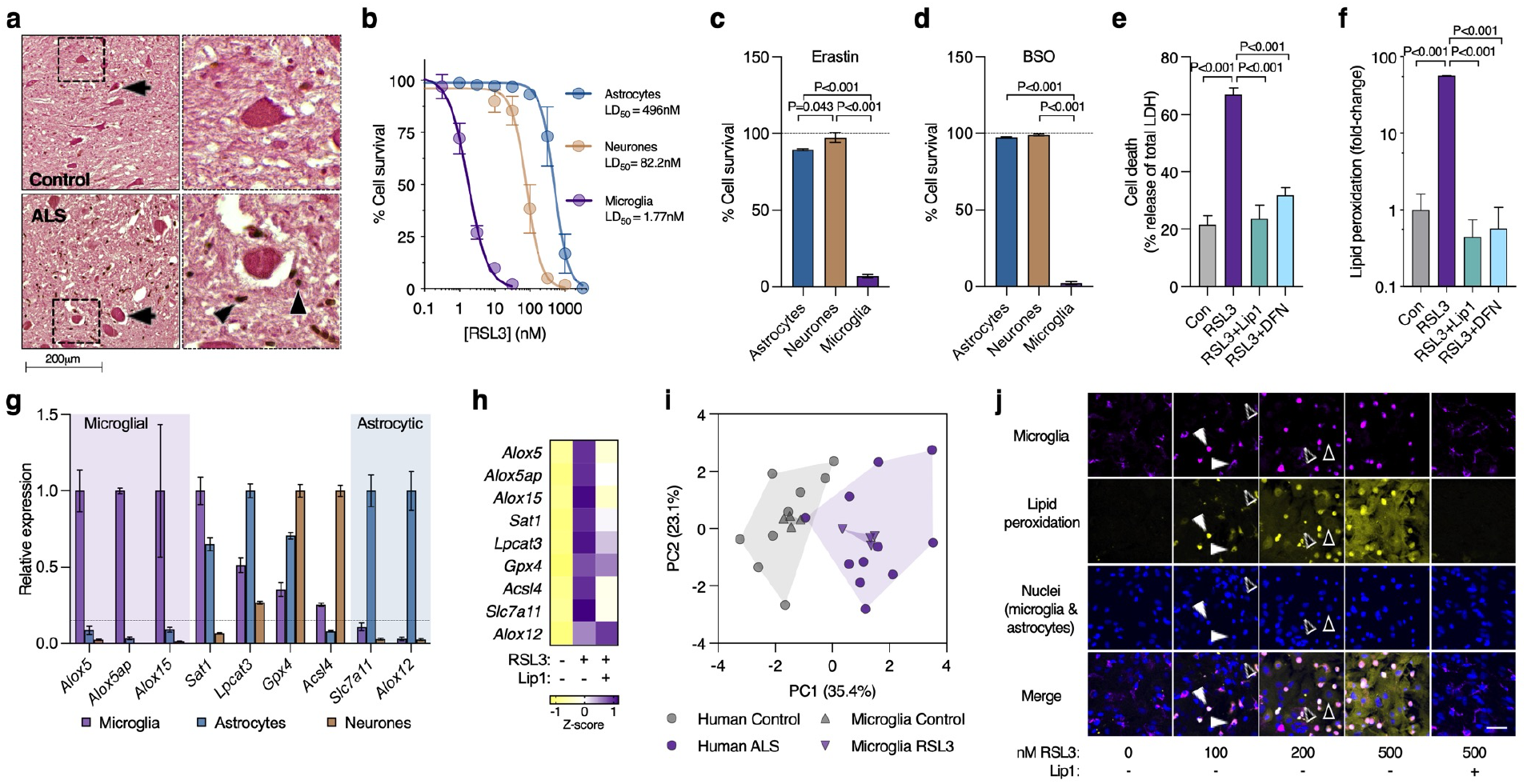
Microglia have heightened sensitivity to ferroptosis. (**a**) Histological assessment of human ALS-affected tissue indicates that iron accumulation in ALS (brown stain; arrow heads) is conspicuously absent from neurones (arrows). (**b-d**) Survival of cultured microglia, astrocytes and neurones exposed to the ferroptosis inducers RSL3, erastin or BSO (n=3-12). (**e-f**) Cytotoxicity and lipid peroxidation in cultured microglia exposed to RSL3 and protection by the inhibitors liproxstatin-1 (Lip1) and deferiprone (DFN) (n=2-3). Lipid peroxidation in **f** measured using oxidised:reduced of the ratiometric fluorophore C11-BODIPY. (**g**) Transcripts associated with ferroptosis in isolated primary murine cultures of microglia, astrocytes and neurones, depicted relative to highest expression. (**h**) Effect of RSL3 on transcripts associated with ferroptosis in cultured microglia, and protection by Lip1 (n=2-4). (**i**) Principal component analysis of ferroptosis genes in response to RSL3 treatment in cultured microglia or ALS in human spinal cord. (**j**) Lipid peroxidation in response to RSL3 and protection with Lip1, in mixed glial cultures (microglia and astrocytes) detected using oxidised:reduced C11-BODIPY (yellow) showing lipid peroxidation is restricted to microglia (white arrowheads) relative to the preponderant astrocytes (black arrowheads). Images derived from **Supplementary Video 1**. Scale bar (**j**) = 50 μm. P values in **c-f** indicate significant differences. Error margins in **b-g** are S.E.M. Individual heatmap values in **h** represent mean. Individual symbols in **i** represent the number of individual control and ALS cases analysed, or independent microglial cultures. Proportion of variance explained by each principal component in **i** is denoted on axes. Data in **i** derived from fold expression change shown in Supplementary **Fig. 2b & 4b**.

Comparison of cultured microglia, astrocytes and neurones revealed that the lipoxygenase genes that were prominently elevated in ALS (*ALOX5, ALOX5AP, ALOX15*; **Fig. 1i; Supplementary Fig. 2b**) were specifically expressed in microglia (**Fig. 2g**) and increased in response to RSL3 treatment in microglia but not astrocytes, an effect mitigated by liproxstatin-1 (**Fig. 2h**; **Supplementary Fig. 4a,b**). The cell-specific expression pattern was corroborated by the BrainRNAseq database (42) (https://www.brainrnaseq.org). Principal component analysis of ferroptosis-related genes revealed RSL3 treatment induced changes in microglia that were similar to those evident in ALS-affected spinal cord (**Fig. 2i**).

Microglial sensitivity to ferroptosis was further characterised using mixed glial cultures comprised of primary murine microglia and astrocytes. Live cell imaging of mixed glial cultures indicated that lipid peroxidation occurs in microglia at a lower dose of RSL3 than in astrocytes (**Fig. 2j; Supplementary Fig. 4c; Supplementary Video 1**). Microglia were also more sensitive than astrocytes to other inducers of ferroptosis (erastin, iron, BSO) in mixed glial cultures (**Supplementary Fig. 4d-h; Supplementary Videos 2 & 3**). Lipid peroxidation was prevented by the ferroptosis inhibitors liproxstatin-1 and deferiprone. Although sufficient to induce microglial lipid peroxidation, neither RSL3 nor erastin plus iron killed microglia under these conditions **(Supplementary Fig. 4i,j)**.

Microglial-mediated activation of neurotoxic astrocytes has been demonstrated using the bacterial endotoxin LPS by sequential transfer of conditioned medium from LPS-activated microglia to astrocytes, then from astrocytes onto cultured neurones (11). Neurotoxicity of conditioned medium from our mixed glial cultures after treating with LPS was consistent with this (**Fig. 3a(i)**). Moreover, applying RSL3 or erastin to the mixed glia instead of LPS also produced conditioned medium that was neurotoxic (**Fig. 3a(i)**), demonstrating that these canonical inducers of ferroptosis, when applied to glial cells, produced a comparable neurotoxic result. The absence of neuronal death from equivalent treatments applied directly to the neurones confirmed the role of glial cells in this model of non-cell autonomous neuronal death (Fig. **3a(v)**). Moreover, mitigation of neurotoxicity from RSL3 or erastin-induced glial conditioned medium by treating the glial cells with liproxstatin-1 supported the role of glial ferroptotic stress as the initiating event (**Fig. 3a(i)**).

**Figure 3.**
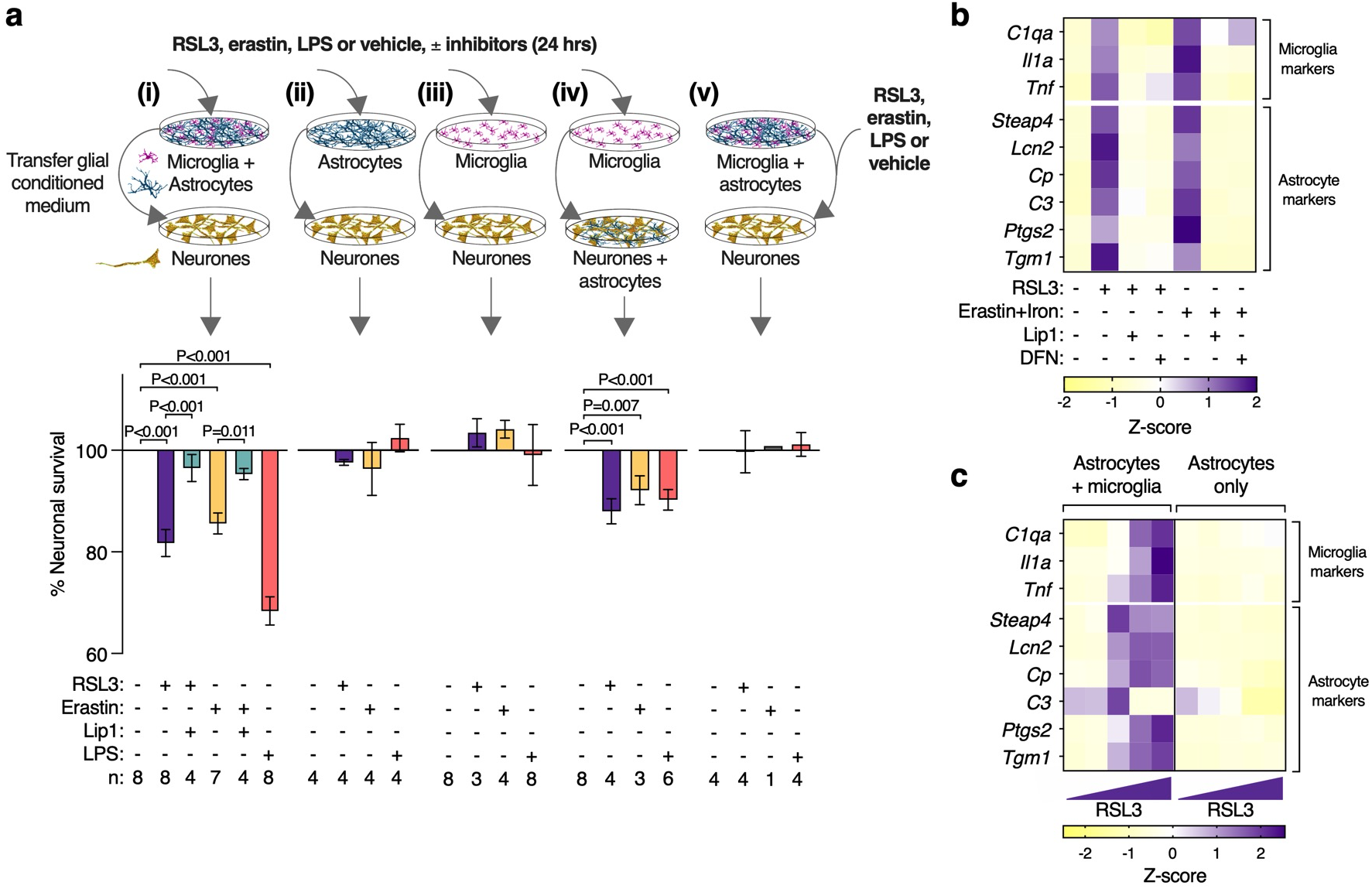
Ferroptotic stress induces neurotoxic glial activation. (**a**) Conditioned medium from mixed glial cultures (**i**) treated with RSL3, erastin or LPS is toxic to cultured neurones. Toxicity is alleviated by liproxstatin-1 (Lip1). Conditioned media from identically treated isolated astrocyte **(ii)** or microglia (**iii**) cultures is not neurotoxic, whereas conditioned medium from isolated microglia is neurotoxic to neurones cultured with astrocytes (**iv**). Adding RSL3, erastin or LPS directly onto neurones is not toxic (**v**). Error margins are S.E.M. Sample size indicated (n) for each condition. P values indicate significant differences. (**b**) Effect of RSL3- or erastin plus iron (as ferric ammonium citrate)-induced ferroptotic stress on expression of genes associated with activation of microglia and astrocytes in mixed glial cultures, and protection by Lip1 or deferiprone (DFN). (**c**) RSL3 induced expression of glial activation genes in mixed glial cultures (astrocytes & microglia) but not astrocytes alone. Individual heatmap values (**b,c**) represent mean (n=3-6). Glial conditioned medium is concentrated using 30 kDa MWCO filters.

Conditioned media in all experiments was fractionated using 30 kDa molecular weight cut-off filters (**Supplementary Fig. 5a**). The concentrated filter retentate increased neurotoxicity (**Supplementary Fig. 5b**), indicating a gain of toxicity in the conditioned medium consistent with neurotoxic astrocyte activation, as opposed to a loss of trophic support. The latter is further supported by the use of sublethal concentrations of RSL3 or erastin excluding glial death as a contributing factor (**Supplementary Fig. 4i,j**). Furthermore, media retained by the filters was neurotoxic whereas media that passed through was not (**Supplementary Fig. 5c**). This indicates neurotoxicity was not instigated by free molecules with a molecular mass <30kDa.

Greater susceptibility of microglia to ferroptotic stimuli than astrocytes and neurones (**Fig. 2b-d,j**) raised the possibility that neurotoxicity of the ferroptosis-induced glial conditioned medium was the result of a cascade of events instigated by a microglial response to ferroptotic stress. Although not involving ferroptosis, the LPS-induced, microglial-mediated activation of neurotoxic astrocytes (11) provides a precedent for such an inflammatory signalling pathway to neuronal death involving an interplay between the two glial cell types. Medium from isolated astrocytes or microglia treated with ferroptosis inducers or LPS was not toxic to neurones (**Fig. 3a(ii,iii)**), whereas microglial conditioned medium was toxic to cultures comprising both neurones and astrocytes (**Fig. 3a(iv)**). This indicates the neurotoxic cascade was initiated in microglia, but astrocytes were required for neurotoxicity.

LPS-treated microglia up-regulate *C1qa, Il1a* and *Tnf*, provoking astrocytes to adopt a neurotoxic phenotype termed A1 with a gene expression signature distinct from alternatively activated astrocytes, termed A2 (11). In our cell culture model in which microglia and astrocytes were grown together, both LPS and RSL3 treatments up-regulated microglial *C1qa, Il1a* and *Tnf* (**Supplementary Fig. 6a**). We analysed expression of a subset of the genes reported for neurotoxic astrocytes (11) and found that RSL3 did not induce a robust A1 phenotype (**Supplementary Fig. 6a**). However, of the genes analysed, a subset was identified whose expression was increased by both RSL3 and LPS (**Supplementary Fig. b,c**), and thus associated with the neurotoxic phenotype. Furthermore, principal component analysis showed that RSL3 (but not LPS) induced changes in these genes that recapitulated changes observed in ALS-affected spinal cord (**Supplementary Fig. 6d)**.

Inducers of ferroptosis applied to mixed glial cultures up-regulated microglial *C1qa, Il1a* and *Tnf*, along with the expression of identified genes associated with neurotoxic glia, and these markers were suppressed by the ferroptosis inhibitors liproxstatin-1 and deferiprone (**Fig. 3b; Supplementary Fig. 6e**). Isolating astrocytes from mixed glial cultures (**Supplementary Fig. 7a-g; Supplementary Video 4**) before treating with ferroptosis inducers abolished the upregulation of genes associated with neurotoxicity (**Fig. 3c; Supplementary Fig. 7h-k**), corroborating the conditioned medium findings (**Fig. 3a**) that the neurotoxic activation of glia by ferroptotic stress requires the presence of microglia.

### Markers of neurotoxic glial activation correlate with ferroptosis in ALS

Complement component 3 (C3) is one of the most prominent molecular markers delineating neurotoxic reactive astrocytes from alternatively activated astrocytes, and is reportedly elevated in neurodegenerative disease (11, 43). Our assessment of C3 protein in human, ALS-afflicted spinal cord corroborates this finding (**Fig. 4a**). Further, our analysis of gene expression changes indicated signatures for A1 and A2 astrocytes, and the subset of genes we identified to be associated with neurotoxic glia are all elevated in ALS-affected spinal cord (**Fig. 4b,c; Supplementary Fig. 2c**). The expression of genes associated with ferroptosis (**Fig. 1i,j**) were positively correlated to those associated with neurotoxic glial activation in human, ALS-affected spinal cord (**Fig. 4d**).

**Figure 4.**
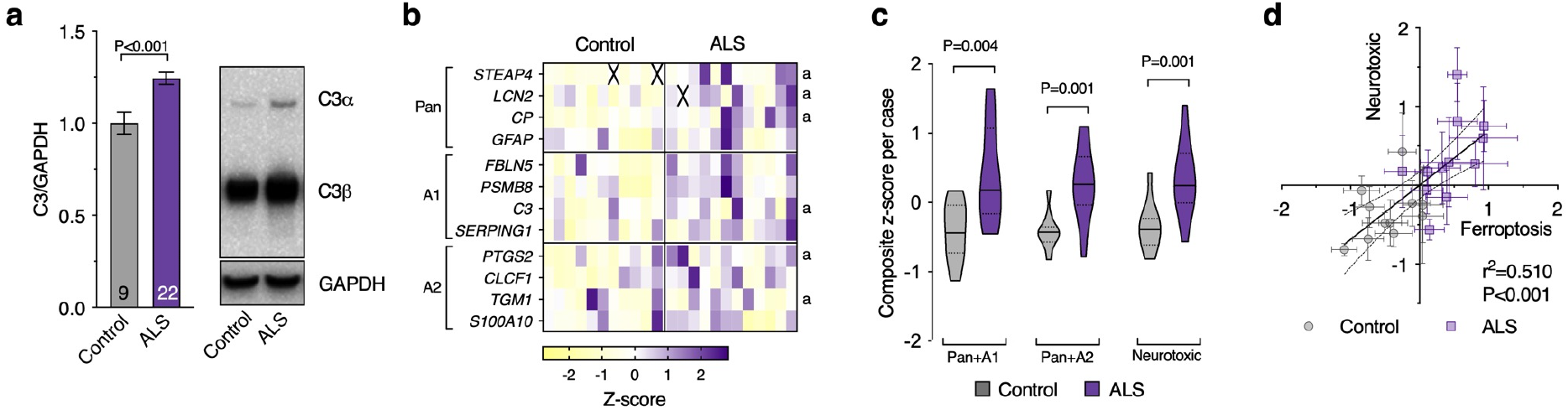
Markers of glial activation and ferroptosis are correlated in human, ALS affected spinal cord. (**a**) Protein levels of neurotoxic astrocyte marker C3 expressed relative to controls. (**b,c**) Transcripts associated with glial activation, highlighting selected markers designated for pan, A1 and A2 activation. Genes associated with neurotoxicity *in vitro* (shown in **Supplementary Fig. 6**) are indicated by ‘a’. Violin plots in **c** represent overall transcript signature for features indicated, derived from heatmap data shown in **b**, including genes associated with neurotoxicity *in vitro* (Neurotoxic). **(d)** Correlation between overall transcript signatures for ferroptosis and glial activation associated with neurotoxicity. Symbols represent composite z-scores for ferroptosis (from **Figure 1j**) and glial activation associated with neurotoxicity (from **c**) for corresponding individual cases. P values show significant differences where indicated (**a,c**) or significance of correlation (**d**). Numbers in bar graph represent the number of individual control and ALS cases analysed. Values in transcript heatmap (**c)** represent z-scores for individual control and ALS cases. Crosses represent excluded samples. Error margins are S.E.M. per mean (**a**) or per case (**d**), or 95% confidence interval of linear regression (dashed lines) in **d**. Solid lines in violin plots represent median, dotted lines represent 25^th^/75^th^ percentiles, truncated at min-max values.

### Cu^II^(atsm) protects against glial ferroptosis and neurotoxic glial activation

The cell and blood-brain barrier permeant Cu^II^(atsm) (**Fig. 5a**) exhibits anti-ferroptotic activity in monocultures of immortalised cell lines, primary neurones and cell-free assays (44, 45). In line with this, we found that Cu^II^(atsm) exhibits anti-ferroptotic activity in glial cells. We found that Cu^II^(atsm) prevented RSL3-induced lipid peroxidation and cytotoxicity in mixed glial cultures at concentrations similar to liproxstatin-1 and ferrostatin-1 (**Fig. 5b; Supplementary Fig. 8a-b**). Cu^II^(atsm) also prevented the cytotoxicity of other inducers of ferroptosis (BSO, erastin, iron; **Supplementary Fig. 8c-d**), RSL3-induced lipid peroxidation and cytotoxicity in isolated microglia **(Supplementary Fig. 8e-h; Supplementary Video 4**), and prevented microglial lipid peroxidation in mixed glial cultures treated with sublethal RSL3 (**Supplementary Fig. 8i-j; Supplementary Video 5**). Furthermore, Cu^II^(atsm) mitigated RSL3-induced up-regulation of ferroptosis-related genes in isolated microglia (**Fig. 5c; Supplementary Fig. 9a**), and gene expression associated with neurotoxic glial activation induced by sublethal RSL3 or erastin in mixed glial cultures **(Fig. 5d; Supplementary Fig. 9b)**. Concordantly, Cu^II^(atsm) suppressed the neurotoxicity of glial conditioned medium induced by RSL3 or erastin (**Fig. 5e**).

**Figure 5.**
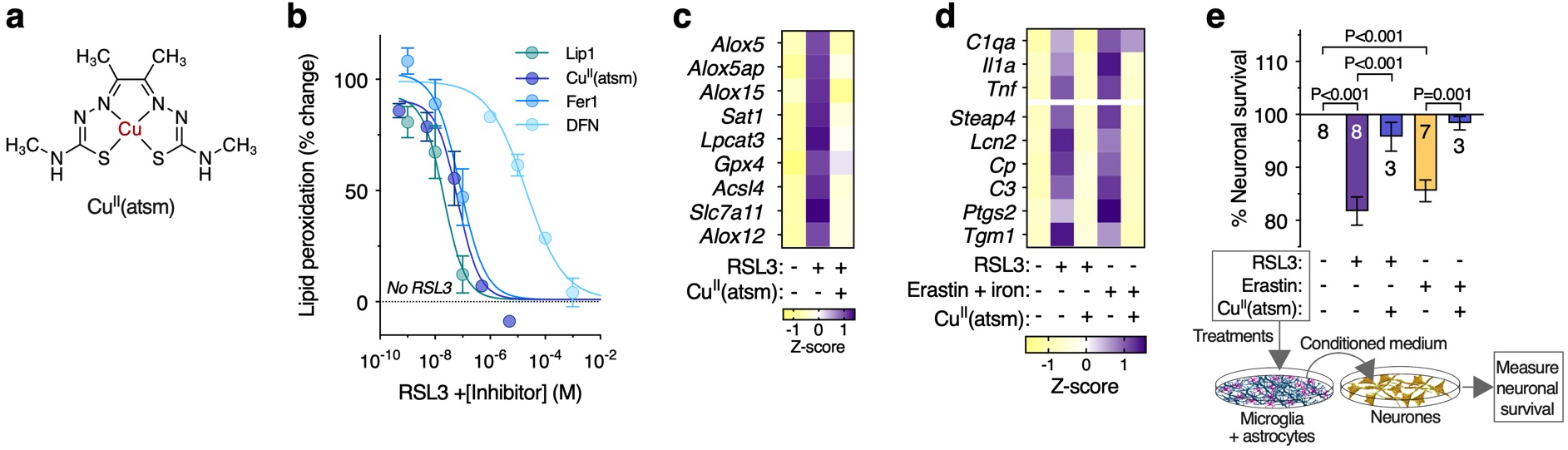
The metallocomplex Cu^II^(atsm) protects against glial ferroptosis and neurotoxic glial activation *in vitro*. (**a**) Chemical structure of Cu^II^(atsm). (**b**) Cu^II^(atsm) prevents RSL3-induced lipid peroxidation in mixed primary cultures of murine microglia and astrocytes, with efficacy similar to the ferroptosis inhibitors liproxstatin-1 (Lip1) and ferrostatin-1 (Fer1). Lipid peroxidation measured as oxidised:reduced C11-BODIPY (n=2-4). (**c**) Cu^II^(atsm) mitigates RSL3-induced expression of genes associated with ferroptosis in microglial cultures (n=3-4). (**d**) Cu^II^(atsm) mitigates ferroptotic stress-induced expression of genes associated with activation of microglia and astrocytes in mixed glial cultures (n=3-6). (**e**) Cu^II^(atsm) inhibits RSL3- and erastin-induced generation of neurotoxic glial conditioned medium, resulting in neuroprotection. Numbers in bars represent number of independent biological replicates. P values indicate significant differences. Error margins in **b,e** are S.E.M. Symbols (**b**) or individual heatmap values (**c,d**) represent mean.

### Pharmacological mitigation of markers of ferroptosis and neurotoxic glial activation associates with disease modification *in vivo*

To further interrogate the relationship between ferroptosis and neurotoxic glial activation, we assessed spinal cord tissue collected from the SOD1^G37R^ mouse model of ALS (22). These analyses consisted of a cohort of animals treated with Cu^II^(atsm) which, supportive of previous studies (23, 46-50), slowed the rate of decline in motor function and resulted in an overall improvement in survival when treatment commenced at the post-symptom onset age of 140 days (**Fig. 6a-d**).

**Figure 6.**
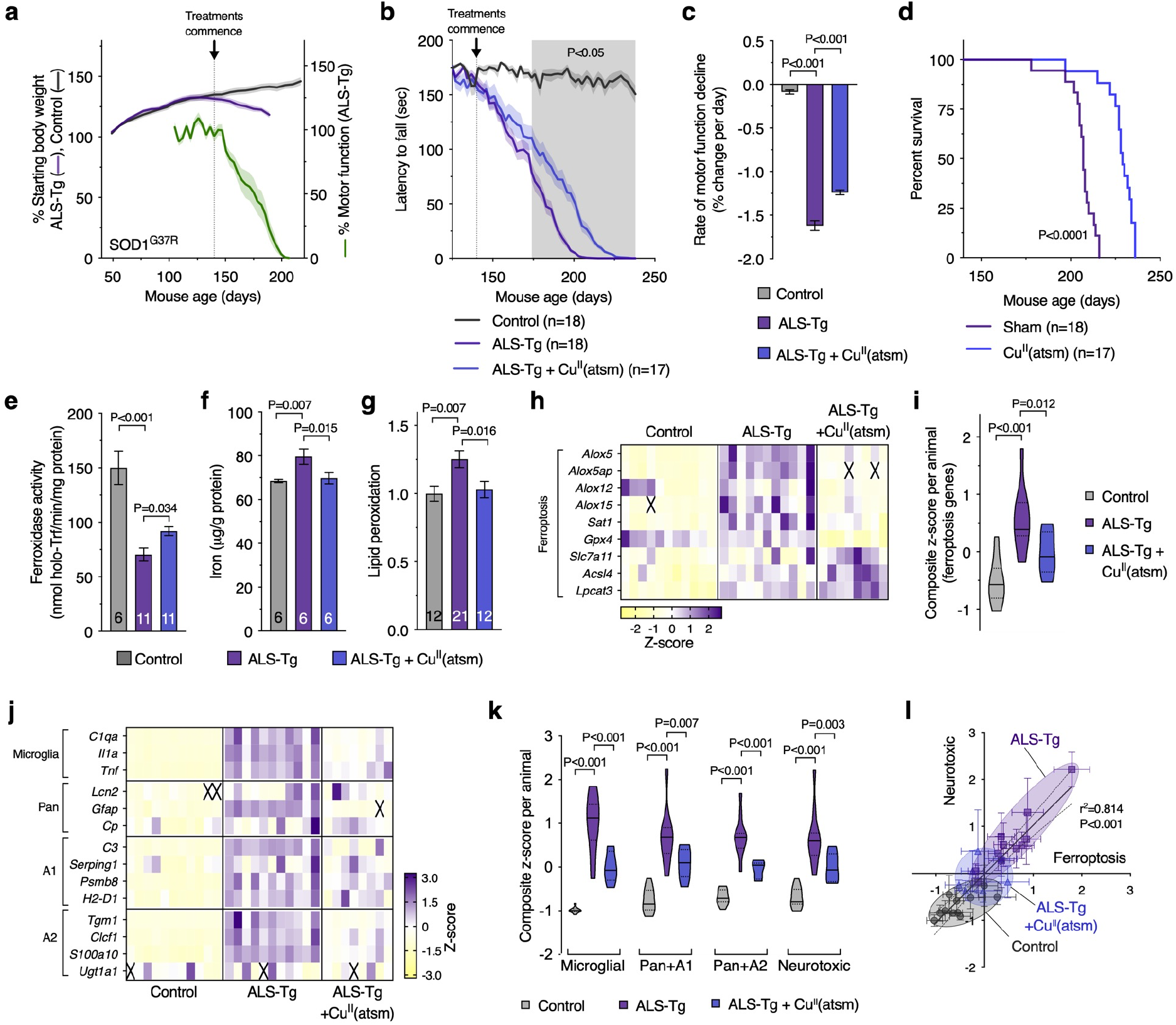
Treatment with Cu^II^(atsm) is protective and mitigates markers of ferroptosis and glial activation in ALS model mice. (**a**) Stage of phenotype progression in transgenic SOD1^G37R^ ALS model mice (ALS-Tg) at which treatment with Cu^II^(atsm) commenced for the present study. Percentage starting body weight for SOD1^G37R^ mice is expressed relative to body weight per animal at 50 days old. Percentage motor function (rotarod assay) is expressed relative to average performance per animal over the period 125-139 days (n=14-21 animals). (**b**) Effect of Cu^II^(atsm) at 30 mg/kg body weight twice daily on motor function of SOD1^G37R^ mice. (**c)** Rate of motor function decline in SOD1^G37R^ mice derived from data in **b**. (**d**) Cu^II^(atsm), administered orally commencing at the post symptom-onset age of 140 days, extends survival of the SOD1^G37R^ mouse model of ALS. (**e-g**) Therapeutic outcomes for Cu^II^(atsm) in the SOD1^G37R^ mice are associated with improved biochemical markers of ferroptosis, including increased ferroxidase activity, decreased iron levels and decreased lipid peroxidation in extracted spinal cord tissue. Lipid peroxidation in **g** measured as oxidised:reduced C11-BODIPY. (**h,i**) Transcript analysis of SOD1G37R mouse spinal cord tissue reveals a molecular profile consistent with ferroptotic stress that is mitigated by treatment with Cu^II^(atsm). (**j,k**) Transcript analysis of SOD1^G37R^ mouse spinal cord tissue reveals a molecular profile consistent with neurotoxic glial activation that is normalised by treatment with Cu^II^(atsm). (**l**) Correlation between overall transcript signatures for ferroptosis and neurotoxic glia (from **i** and **k**). Symbols represent individual mice. Numbers in each bar graph represent the number of individual mice analysed. Crosses in heatmaps represent excluded samples. Individual values in transcript heatmaps (**h,j**) represent z-scores for individual animals. Violin plots represent overall transcript signature for ferroptosis (**i**) or activation of microglia and astrocytes (**k**), derived from heatmap data shown in **h** and **j**. Error margins (dashed lines) in correlation plot (**l**) are 95% confidence interval of linear regression, and S.E.M. in line plots (**a,b**), bar graphs (**c,e-g**) and overall transcript signatures of individual mice **(l**). Solid lines in violin plots **(i,k)** represent median, dotted lines represent 25^th^/75^th^ percentiles, truncated at min-max values. P values indicate significant differences between groups (**c-g,i,k**), significance of correlation (l), or significant difference between SOD1^G37R^ mice treated with or without Cu^II^(atsm) (shaded area, **b**).

Analysis of spinal cord tissue collected from animals at the mid-symptom age of 175-195 days revealed loss of ferroxidase activity, accumulation of iron, and increased lipid peroxidation, with all three mitigated by treatment with Cu^II^(atsm) **(Fig. 6e-g**). A gene expression signature consistent with ferroptosis was also evident in the mouse model (**Fig. 6h,i; Supplementary Fig. 10a**). These data show that features of ferroptosis evident in human, sporadic cases of ALS are reproduced in this mutant SOD1 model of the disease. Treatment with Cu^II^(atsm) attenuated the ferroptosis gene signature, consistent with the *in vitro* anti-ferroptosis activity of Cu^II^(atsm) recently reported (44) and as we found in cultured glial cells (**Fig. 5**).

Molecular markers of neurotoxic glial activation were also elevated in the SOD1^G37R^ mouse model (**Fig. 6j,k; Supplementary Fig. 10b**), consistent with data reported for the SOD1^G93A^ model (43). These changes correlated with the expression signature for ferroptosis (**Fig. 6l**), thus reproducing the association between ferroptosis and neurotoxic glial activation observed in human, sporadic ALS-affected spinal cord (**Fig. 4d**). Consistent with ferroptosis inducing glial activation *in vivo*, the molecular markers of glial activation in the SOD1^G37R^ mouse model were suppressed by treatment with Cu^II^(atsm) (**Fig. 6j-l; Supplementary Fig. 10b**). Principal component analysis revealed that changes in expression of genes related to ferroptosis or to neurotoxic glial activation were similar between glial cells treated with RSL3, sporadic ALS-affected tissue, and the SOD1^G37R^ mouse model (**Supplementary Fig. 10c,d**).

## Discussion

By demonstrating that ferroptotic stress can trigger neurotoxic glial activation, data presented herein provide new insight to how glial cells can cause non-cell autonomous neuronal death in neurodegenerative disease. Our proposed pathway from ferroptotic stress to neuronal death is supported by results from primary glial and neuronal cells grown in culture (**Fig. 2 & 3**), its involvement in human disease is indicated by results from ALS-affected spinal cord tissue (**Fig. 1 & 4**), and the potential for therapeutic mitigation is supported by *in vitro* and *in vivo* data from experiments involving the CNS permeant ferroptosis inhibitor Cu^II^(atsm) (**Fig. 5 & 6**).

As there are no specific markers of ferroptosis, its identification in tissue samples is established by the accumulation of consistent evidence across diverse markers. To date, different lines of evidence implicate ferroptosis in prominent neurodegenerative diseases including ALS. Here, our multifaceted analyses spanning atomic, molecular and biochemical indications provide the first comprehensive evidence indicating ferroptosis is a salient feature of sporadic ALS-affected spinal cord (**Fig. 1**). Moreover, these changes are recapitulated in a robust animal model of ALS (**Fig. 6**), supporting the veracity of this model to investigate ferroptosis in ALS.

The current common perception of ferroptosis is that it is an autonomously initiated cell suicide event (51). Ferroptosis inducers or inhibitors have been used to interrogate the role of ferroptosis in *in vivo* models of neural disease, including Huntington’s disease, Parkinson’s disease, ALS, ischemic stroke and intracerebral haemorrhage (52-57). These studies, however, provide little insight into the cellular consequences of ferroptosis and the results are generally interpreted as evidence for direct neuronal ferroptosis. Our findings necessitate a re-evaluation of these interpretations. Here we demonstrate a novel pathophysiological role for sublethal ferroptotic stress culminating in neuronal death in a non-cell autonomous manner. This proposition is supported by our findings that microglia – the resident immune cells of the CNS – are more sensitive to ferroptosis than astrocytes or neurones *in vitro* (**Fig. 2**), an observation in line with recent gene expression data (58). When ferroptotic stress insufficient to induce cell death is applied to microglia, it triggers an inflammatory response that induces a neurotoxic glial phenotype (**Fig. 3**). This neurotoxicity only occurs in the presence of astrocytes. Furthermore, we find corroborative evidence for the presence of neurotoxic glia in sporadic ALS-affected tissue (**Fig. 4**) and the SOD1^G37R^ mouse model of ALS (**Fig. 6**), where the expression signatures for ferroptosis genes and neurotoxic glia are correlated in both. Finally we observe that the gene expression signatures for ferroptosis and neurotoxic glial activation are highly similar for ferroptosis-stimulated glia *in vitro*, for sporadic ALS-affected tissue, and for ALS model mice (**Supplementary Fig. 10c,d**). Thus, our results indicate that neuronal demise may not be the result of overt neuronal ferroptosis, but rather an inflammatory signalling pathway instigated by microglial ferroptotic stress in the absence of overt ferroptotic cell death. Our data indicate that ferroptotic stress preferentially influences microglia over other CNS cells. This is supported by multiple lines of evidence: 1) Ferroptotic stress that causes lipid peroxidation, induction of ferroptosis-related genes and an inflammatory response in microglia has no discernable impact on neurones or astrocytes; 2) of the ferroptosis-related genes induced by ferroptosis, many coding for lipoxygenases are specifically expressed by microglia (**Fig. 2g**) and are elevated in sporadic ALS-affected tissue (**Fig. 1**) and ALS model mice (**Fig. 6**), but not in ferroptosis-exposed astrocytes (**Supplementary Fig. 4a**); 3) lipoxygenases are involved in intitiation of ferroptosis (40) and their specific expression by microglia in the CNS may contribute to the greater sensitivity of microglia to ferroptosis (**Fig. 2**); and 4) microglial gene expression changes identified by RNAseq in response to ferroptotic stimuli are evident in human ALS-affected spinal cord (58). Together, these data indicate that the emergence of ferroptotic stimuli *in vivo*, as evident in ALS, causes preferential inflammatory activation of microglia that induces neurotoxic activation of astrocytes and neuronal death.

The relevance of microglial activation of neurotoxic astrocytes in neurodegenerative disease was recently substantiated where genetic deletion of *Il1a, Tnf* and *C1qa* produced strong neuroprotective and disease-modifying outcomes in a SOD1^G93A^ mouse model of ALS, including a 54% extension to survival (43). Other disease-relevant triggers of neurotoxic glial activation include release of damaged mitochondria from microglia in cell culture models of ALS, Huntington’s disease and Alzheimer’s disease (59), and fibrils of α-synuclein in Parkinson’s disease (60). These studies support the assertion that inflammatory activation of microglia causes neurotoxic glial activation in neurodegeneration.

The role of ferroptosis in ALS is supported by the overexpression of GPX4 delaying symptom onset and improving survival of SOD1^G93A^ ALS model mice (56). Our results lead us to propose that the ferroptotic stress-induced neurotoxic glia pathway is a treatable feature of ALS. *In vitro*, canonical ferroptosis inhibitors blocked upregulation of ferroptosis genes in microglia, prevented upregulation of neurotoxic glial activation genes and inhibited the neurotoxicity of ferroptosis-induced glial conditioned medium (**Fig. 2, 3**). These results provide further evidence for the causal relationship between ferroptosis and neurotoxic glial activation. Furthermore, these effects were recapitulated by treatment with Cu^II^(atsm) – a compound that we and others confirm has direct anti-ferroptotic activity (**Fig. 5**) (44, 45). Cu^II^(atsm) also suppressed markers of both ferroptosis and neurotoxic glia evident in the ALS model mice (**Fig. 6**). Together, these data demonstrate a causal relationship between microglial ferroptosis, neurotoxic astrocyte activation and neuronal death *in vitro* that is strongly corroborated *in vivo*.

## Conclusions

Results presented herein provide the first evidence for reactive microglia responding to ferroptotic stress causing non-cell autonomous neuronal death. The neuropathology of diverse neurodegenerative conditions features reactive gliosis (61). These diseases also exhibit signs of ferroptosis (including iron accumulation, glutathione depletion, and lipid peroxidation (62-65)) and the involvement of neurotoxic astrocytes is implicated (11). Accordingly, the ferroptosis-neurotoxic glia pathway we describe here and illustrate in the context of ALS may contribute to neuronal death in other neurodegenerative diseases. By demonstrating that sublethal ferroptotic stress in microglia is a non-cell autonomous trigger of neuronal death, our discovery provides new understanding of how neurones die in neurodegenerative disease. These results also highlight that therapeutic strategies for neurodegenerative disease that target ferroptosis should not focus solely on neuronal events and need to address glial ferroptosis.

## Abbreviations used

ALS: Amyotrophic lateral sclerosis
BSO: Buthionine sulpoxamine
CNS: Central nervous system
C3: Complement component 3
DFN: Deferiprone
DMEM: Dulbecco’s modified Eagle’s medium
FAC: ferric ammonium citrate
FBS: Foetal bovine serum
GPX4: Glutathione peroxidase 4
ICP-MS: Inductively coupled plasma-mass spectrometry
IMDM: Iscove modified Dulbecco media
LDH: Lactate dehydrogenase
Lip1: Liproxstatin-1
LPS: Lipopolysaccharide
MTT: 3-(4,5-Dimethylthiazol-2-yl)-2,5-diphenyltetrazolium bromide
PC: L-α-Phosphatidylcholine
RCF: Relative centrifugal force
RSL3: (1*S*,3*R*)-Methyl-2-(2-chloroacetyl)-2,3,4,9-tetrahydro-1-[4-(methoxycarbonyl)phenyl]-1*H*-pyrido[3,4-b]indole-3-carboxylic acid
RT-PCR: Reverse transcription polymerase chain reaction
SDS-PAGE: Sodium dodecyl sulphate–polyacrylamide gel electrophoresis
SOD1: Superoxide dismutase 1
SSV: Standard suspension vehicle
TBS: Tris-buffered saline

## Declarations

### Ethics approval

The use of mice for generating primary cell cultures and assessing the effect of Cu^II^(atsm) was approved by a University of Melbourne Animal Experimentation Committee (Projects 1613931, 1814576, 1915081) and complied with National Health and Medical Research Council guidelines. Procedures involving post-mortem human tissue were approved by a University of Melbourne Human Ethics Committee (Project ID 1238124) or University of Sydney Human Ethics Committee (Project ID 2015/202).

## Ackowledgements

Human tissue samples were obtained from the Victorian Brain Bank (Florey Institute of Neuroscience and Mental Health, the University of Melbourne, Australia), the MS Society Tissue Bank (Wolfson Neuroscience Laboratories, Imperial College London, United Kingdom), MRC London Neurodegenerative Diseases Brain Bank (King’s College, London, UK), the University of Maryland Brain and Tissue Bank, a biorepository of the NIH NeuroBioBank (Maryland, USA), and the Sydney Brain Bank. Dr Antonella Roveri (University of Padova, Italy) provided invaluable advice on measuring GPX4 activity in tissue extracts. All live cell imaging was conducted at the University of Melbourne Biological Optical Microscopy Platform with expert advice from Dr Ellie Hyun-Jung Cho. Histological assessment of human spinal cord was performed at the Histology and Neuropathology Facility (Florey Institute of Neuroscience and Mental Health, the University of Melbourne, Australia) by Dr Ian Birchall.

## Funding

This research was supported by funding from Motor Neurone Disease Research Australia (Beryl Bayley Fellowship; Betty Laidlaw MND Research Project; Jenny Barr Smith MND Research Project), FightMND (Translational Research Grant), the Australian Research Council, the National Health and Medical Research Council (Projects 1061550, 1054325, 1181864), and the University of Melbourne. JBH was a recipient of an Australian Postgraduate Award and the Nancy Frances Curry Scholarship. JRL was a recipient of an NHMRC Early Career Fellowship. KK was recipient of a Sigrid Jusélius Foundation Fellowship. PSD was a recipient of an Australian Research Council Future Fellowship (FT2). DJH was a recipient of an NHMRC Industry Career Development Fellowship (CDF1) in partnership with Agilent Technologies.

ARW was a recipient of an NHMRC Senior Research Fellowship. PJC was a recipient of an NHMRC Career Development Fellowship (CDF2, 1084927). The contribution of KLD and BGT to this study was supported by ForeFront, a large collaborative research group dedicated to the study of neurodegenerative diseases and funded by the National Health and Medical Research Council.

## Author contributions

JRL, JBWH, KK, AIB and PJC contributed to conception and design of the study, acquisition and analysis of data, and drafting the manuscript. SN, LEM, DJH, BP, BGT, KLD, SWM, SA, BRR, JSB, CAM, ARW and PSD contributed to acquisition and analysis of data.

## Competing interests

Collaborative Medicinal Development LLC has licensed intellectual property related to this subject from the University of Melbourne where the inventors include ARW and PSD. AIB is a shareholder in Alterity Ltd, Cogstate Ltd, Brighton Biotech LLC, Grunbiotics Pty Ltd, Eucalyptus Pty Ltd, and Mesoblast Ltd. He is a paid consultant for Collaborative Medicinal Development LLC and has a profit share interest in Collaborative Medicinal Development Pty Ltd. PJC and JSB are unpaid consultants for Collaborative Medicinal Development LLC.

## Availability of data

The datasets used and/or analysed during the current study are available from the corresponding author on reasonable request.

## Author information

Jeffrey R. Liddel, James B.W. Hilton, and Kai Kysenius contributed equally to this work.

