## Supplementary Material for "Microglial ferroptotic stress causes non-cell autonomous neuronal death"

Supplementary Fig. 1. Iron content of human, ALS-affected spinal cord.
Supplementary Fig. 2. Gene expression changes in human, ALS-affected spinal cord.
Supplementary Fig. 3. GPX4 in human, ALS-affected spinal cord.
Supplementary Fig. 4. Microglial ferroptosis.
Supplementary Fig. 5. Retention of neurotoxic factor(s) by 30 kDa MWCO filter.
Supplementary Fig. 6. Gene expression changes used to monitor neurotoxic glial activation in response to ferroptotic stress in glial cells.
Supplementary Fig. 7. Isolating astrocytes from mixed glial cultures and gene expression changes in response to treatment with RSL3 or RSL3 plus iron.
Supplementary Fig. 8. Protective activity of $Cu^{II}$(atsm) *in vitro*.
Supplementary Fig. 9. Gene expression changes in mixed glial cultures treated with inducers of ferroptosis and the metallocomplex $Cu^{II}$(atsm).
Supplementary Fig. 10. Gene expression changes in $SOD1^{G37R}$ mice compared to human ALS-affected spinal cord and cultured glia.

Supplementary Video 1. Live cell imaging of lipid peroxidation in mixed glial cultures treated with the ferroptosis inducer RSL3.
Supplementary Video 2. Live cell imaging of lipid peroxidation in mixed glial cultures treated with various ferroptotic stimuli (RSL3, erastin+iron, and iron).
Supplementary Video 3. Live cell imaging of lipid peroxidation in mixed glial cultures treated with BSO.
Supplementary Video 4. Live cell imaging of showing mitigation of lipid peroxidation by the ferroptosis inhibitors, liproxstatin-1 (Lip1), $Cu^{II}$(atsm) and deferiprone (DFN) in isolated microglia treated with RSL3.
Supplementary Video 5. Live cell imaging of showing mitigation of lipid peroxidation by the ferroptosis inhibitor $Cu^{II}$(atsm) in mixed glial cultures treated with RSL3.

**a**

| Sample code | Disease group | Site of onset | Sex | Age | PMI (hrs) |
|---|---|---|---|---|---|
| C1 | Control | N/A | Male | 52 | 33 |
| C2 | Control | N/A | Male | 64 | 68 |
| C3 | Control | N/A | Male | 86 | 71 |
| C4 | Control | N/A | Female | 89 | 6 |
| C5 | Control | N/A | Male | 76 | 50 |
| C6 | Control | N/A | Male | 85 | 68 |
| C7 | Control | N/A | Female | 78 | 23 |
| C8 | Control | N/A | Female | 89 | 22 |
| C9 | Control | N/A | Male | 68 | 30 |
| C10 | Control | N/A | Male | 90 | 12 |
| C11 | Control | N/A | Male | 66 | 54 |
| C12 | Control | N/A | Female | 89 | 13 |
| C13 | Control | N/A | Female | 51 | 33 |
| C14 | Control | N/A | Male | 63 | 23 |
| C15 | Control | N/A | Female | 73 | 27 |
| C16 | Control | N/A | Male | 65 | 25 |
| C17 | Control | N/A | Male | 50 | 24 |
| C18 | Control | N/A | Male | 48 | 22 |
| C19 | Control | N/A | Female | 49 | 26 |
| C20 | Control | N/A | Male | 55 | 23 |
| C21 | Control | N/A | Male | 49 | 22 |
| C22 | Control | N/A | Male | 50 | 22 |
| ALS1 | Sporadic ALS | Limb - lower | Female | 56 | 45 |
| ALS2 | Sporadic ALS | Insuf clinical | Female | 80 | 23 |
| ALS3 | Sporadic ALS | Bulbar | Male | 54 | 18 |
| ALS4 | Sporadic ALS | Limb - upper & lower | Male | 73 | 21 |
| ALS5 | Sporadic ALS | Limb - lower (left) | Male | 65 | 45 |
| ALS6 | Sporadic ALS | Insuf clinical | Male | 72 | 67 |
| ALS7 | Sporadic ALS | Limb - upper & lower | Male | 65 | 14 |
| ALS8 | Sporadic ALS | Limb - upper & lower | Male | 79 | 31 |
| ALS9 | Sporadic ALS | Limb - upper & lower, & chest | Male | 64 | 14 |
| ALS10 | Sporadic ALS | Limb - lower | Female | 77 | 25 |
| ALS11 | Sporadic ALS | Limb - lower | Female | 74 | 7 |
| ALS12 | Sporadic ALS | Insuf clinical | Female | 55 | 25 |
| ALS13 | ALS* | Limb - lower | Male | 80 | 25 |
| ALS14 | Sporadic ALS | Bulbar | Male | 83 | 36 |
| ALS15 | Sporadic ALS | Limb - upper & lower | Male | 58 | 22 |
| ALS16 | Sporadic ALS | Limb - upper | Male | 55 | 56 |
| ALS17 | Sporadic ALS | Limb - lower | Female | 59 | 20 |
| ALS18 | Sporadic ALS | Limb - upper | Male | 54 | 55 |
| ALS19 | Sporadic ALS | Insuf clinical | Female | 55 | 28 |
| ALS20 | Sporadic ALS | Bulbar | Female | 59 | 58 |
| ALS21 | Sporadic ALS | Limb - lower | Male | 71 | 30 |
| ALS22 | Sporadic ALS | Limb - upper (right) | Male | 69 | 36 |
| ALS23 | Sporadic ALS | Limb | Male | 65 | 22 |
| ALS24 | Sporadic ALS | Limb - upper & lower | Male | 71 | 22 |
| ALS25 | Sporadic ALS | Limb - upper | Male | 68 | 66 |
| ALS26 | Sporadic ALS | Limb - upper (right) | Male | 67 | 61 |
| ALS27 | Sporadic ALS | Limb - lower | Male | 62 | 31 |
| ALS28 | Sporadic ALS | Limb - upper & lower | Male | 78 | 14 |
| ALS29 | Familial ALS# | Limb - upper & lower (right) | Female | 69 | 45 |
| ALS30 | Sporadic ALS | Bulbar | Female | 70 | 17 |
| ALS31 | Sporadic ALS | Limb - lower | Female | 74 | 7 |
| ALS32 | Sporadic ALS | Limb - lower | Female | 75 | 9 |
| ALS33 | Sporadic ALS | Bulbar | Female | 86 | 64 |
| ALS34 | Familial ALS - SOD1 | Bulbar | Male | 47 | 14 |
| ALS35 | Familial ALS - SOD1 | Bulbar | Female | 70 | 56 |
| ALS36 | Familial ALS - SOD1 | Limb - upper | Female | 46 | 5 |
| ALS37 | Familial ALS - C9ORF72 | Bulbar | Male | 64 | 68 |
| ALS38 | Familial ALS# | Limb - lower (left) | Male | 65 | 7 |
| ALS39 | Familial ALS - C9ORF72 | Bulbar | Male | 62 | 20 |
| ALS40 | Familial ALS - C9ORF72 | Limb - upper (right) | Male | 55 | 24 |
| ALS41 | Sporadic ALS | Limb - upper | Female | 70 | 15 |
| ALS42 | Sporadic ALS | Limb - lower | Female | 50 | 56 |
| ALS43 | Sporadic ALS | Limb - lower | Female | 68 | 17 |
| ALS44 | Sporadic ALS | Limb - upper and lower | Female | 54 | 12 |
| ALS45 | Sporadic ALS | Limb - lower | Male | 56 | 22 |
| ALS46 | Sporadic ALS | Limb - upper and lower | Male | 68 | 4 |
| ALS47 | Sporadic ALS | Limb - lower (right) | Female | 57 | 20 |
| ALS48 | Sporadic ALS | Limb - lower | Female | 70 | 29 |

*Adopted, no known genetic family history; #Mutation not known; Sporadic ALS, no known family history; C13-C22 and ALS34-ALS49 only used for glutathione analyses; Insuf clinical: Insufficient clinical information available; PMI: Post-mortem interval.

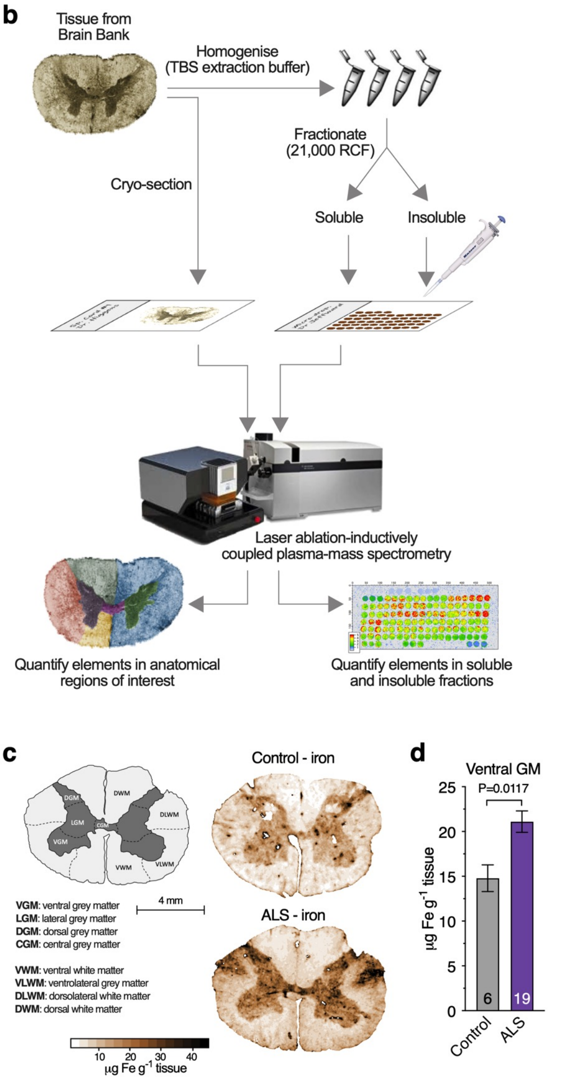


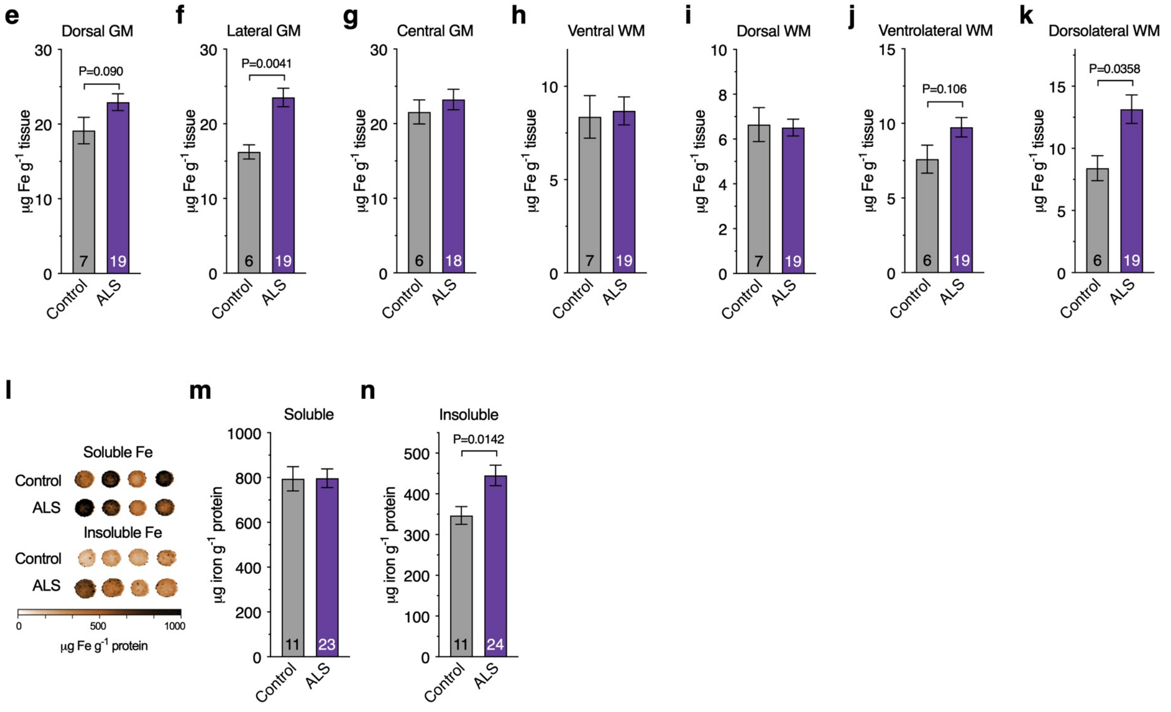


**Supplementary Fig. 1. Iron content of human, ALS-affected spinal cord.**

**(a)** Clinical details for study ALS and control cases. **(b)** Procedure for direct elemental quantitation of iron in human spinal cord samples. Cryo-sectioning followed directly by laser ablation inductively coupled plasma-mass spectrometry (LA-ICP-MS) enables *in situ* quantitation. Prior homogenisation in TBS-based extraction buffer and centrifugation enables quantitation of soluble and insoluble partitioning. **(c)** Anatomical map illustrating regions of interest for *in situ* quantitation of iron and representative heatmaps for iron in spinal cord sections (reproduced from **Figure 1a**). **(d-k)** Iron concentration in indicated anatomical regions of interest. **(l)** Representative images of iron in "micro-droplets" of TBS-soluble and -insoluble fractions of human spinal cord homogenates. **(m,n)** Concentration of iron in TBS-soluble and -insoluble fractions of human spinal cord homogenates. Numbers in bar graphs represent number of individual control and ALS cases analysed. Error margins are S.E.M. and P values indicate significant differences.

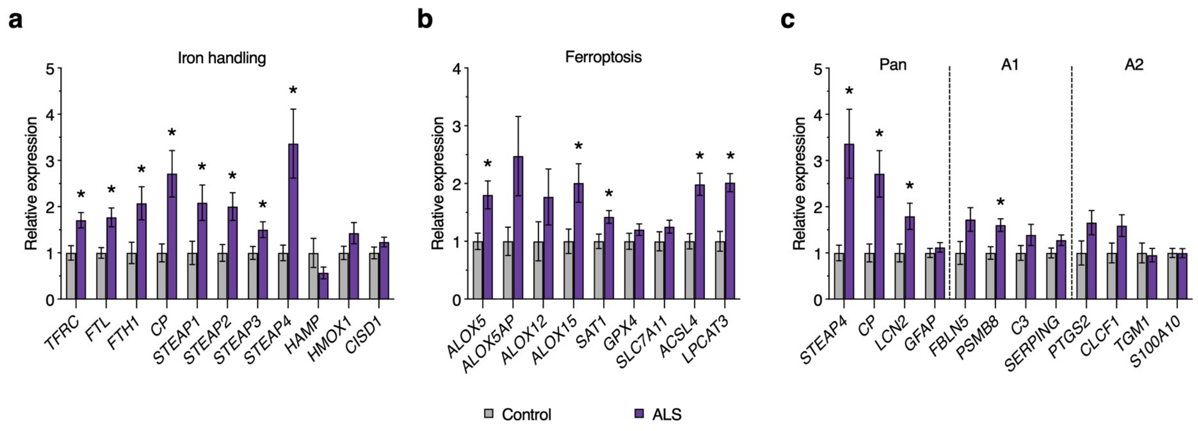


**Supplementary Fig. 2. Gene expression changes in human, ALS-affected spinal cord.**

Relative expression changes for genes associated with **(a)** iron-handling mechanisms, **(b)** ferroptosis, and **(c)** neurotoxic glial activation, with the latter highlighting selected markers designated for pan, A1 and A2 activation. These data are presented in heatmaps shown in **Figures 1d, 1i** and **4b** respectively. Asterisks illustrate significant differences ($P<0.05$) between ALS and respective control. Error margins are S.E.M. (n=9-12).

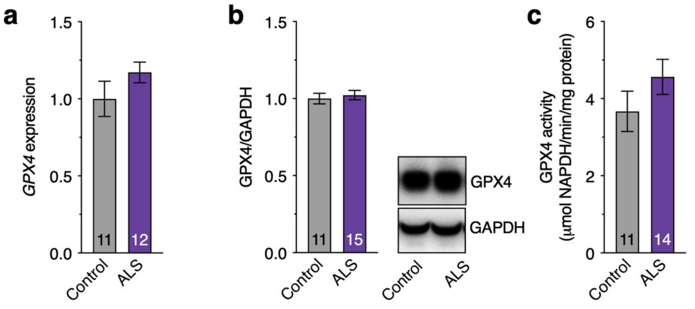


**Supplementary Fig. 3. GPX4 in human, ALS-affected spinal cord.**

**(a)** *GPX4* gene expression in human spinal cord tissue determined by qPCR. **(b)** GPX4 protein levels in human spinal cord tissue determined by western blot. **(c)** GPX4 enzyme activity in human spinal cord tissue determined by rate of RSL3-sensitive, phosphatidylcholine hydroperoxide dependent NADPH consumption. Numbers in bar graphs represent number of individual control and ALS cases assessed. Error margins are S.E.M.

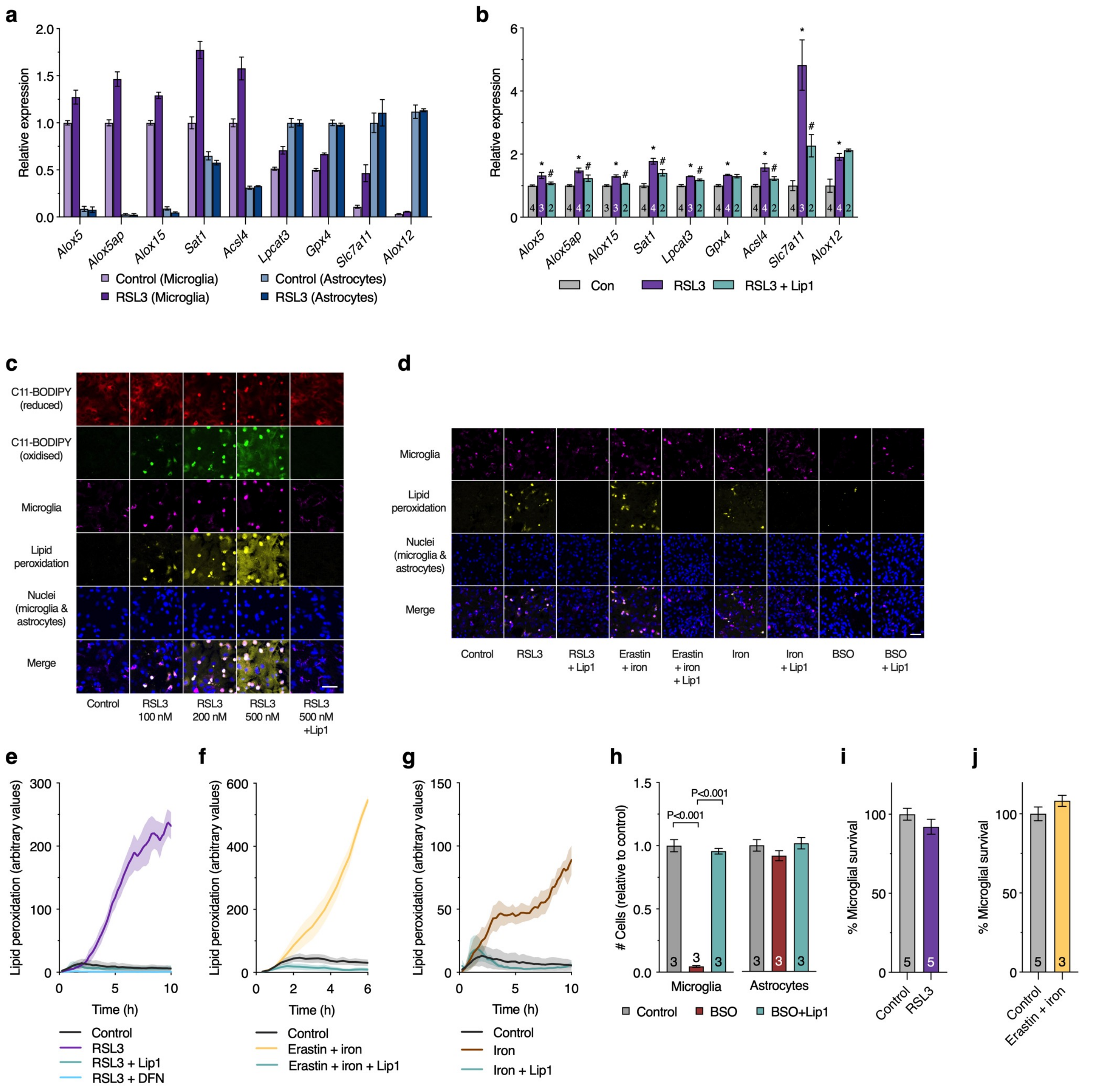


**Supplementary Fig. 4. Microglial ferroptosis.**

**(a)** Transcripts associated with ferroptosis in isolated primary murine cultures of microglia or astrocytes after treating with or without RSL3, normalised to highest control expression (n=3-4). **(b)** Changes in transcripts associated with ferroptosis in cultured microglia and protective effect of Lip1. These data are presented in heatmap shown in **Figure 2h**. **(c)** Lipid peroxidation (oxidized:reduced, yellow) assessed using the ratiometric lipid peroxidation probe C11-BODIPY in mixed glial cultures treated with indicated concentrations of RSL3 showing astrocytic lipid peroxidation only at higher RSL3 concentrations and mitigation with liproxstatin-1 (Lip1; visualised in **Supplementary Video 1**). **(d)** Lipid peroxidation in mixed glial cultures treated with alternate inducers of ferroptosis and the protective effect of Lip1 (visualised in **Supplementary Videos 2 & 3**). **(e-g)** Time dependent changes in microglial lipid peroxidation assessed in mixed glial cultures treated with RSL3, erastin and iron, or iron alone, and mitigation with ferroptosis inhibitors Lip1 or deferiprone (DFN; visualised in **Supplementary Video 2**; n=3-4 for cells treated with ferroptosis inducers and controls; n=1-4 for cells also treated with Lip1 or DFN). **(h)** Number of surviving microglia and astrocytes in mixed glial cultures treated with BSO and the protective activity of Lip1 (visualised in **Supplementary Video 3**). **(i-j)** Microglial survival in mixed glial cultures treated with RSL3 or erastin and iron determined by transcript analysis of microglial-specific genes. Numbers in bars represent number of independent experimental replicates. Iron in **d**, **f**, **g**, **j** is ferric ammonium citrate. Error margins are S.E.M. P values indicate significant differences (**h**). Significant differences between control and RSL3 are indicated by asterisks and between RSL3 treated with or without Lip1 by hash symbols (**b**). Scale bar (**c**,**d**) = 50 µm.

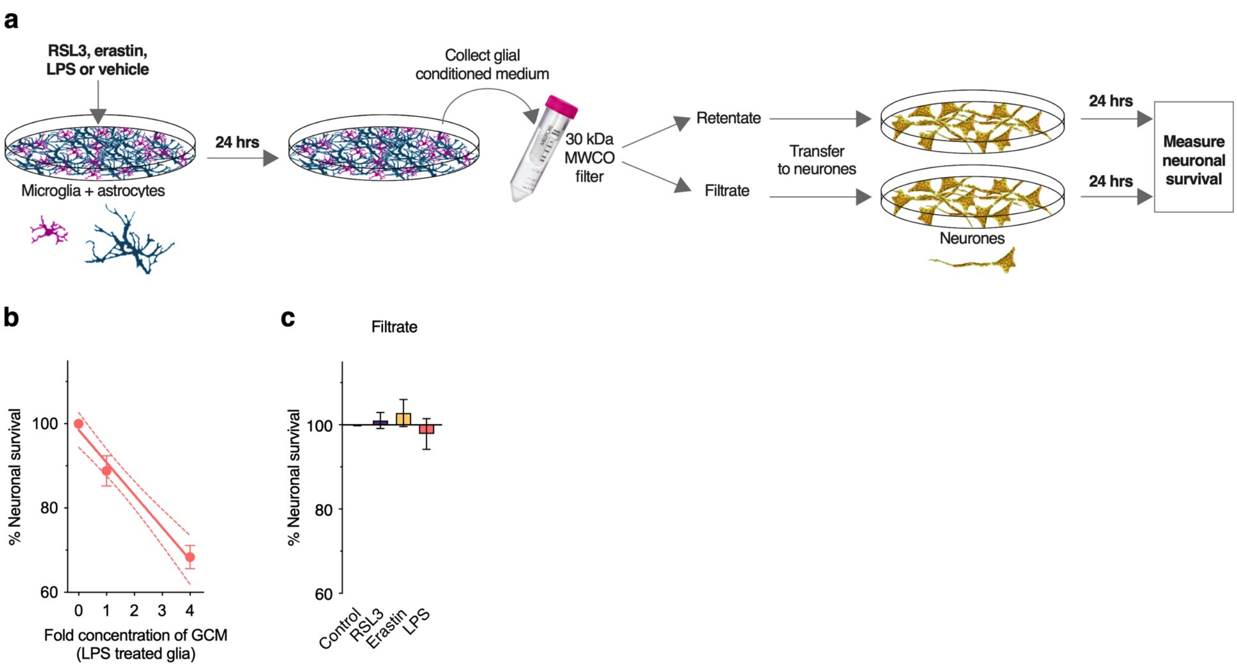


**Supplementary Fig. 5. Retention of neurotoxic factor(s) by 30 kDa MWCO filter.**

**(a)** Procedure for exposing neurones to glial conditioned medium. RSL3, erastin, LPS or vehicle control is added to mixed glial cells (microglia and astrocytes; depicted), isolated astrocytes or isolated microglia for 24 hrs. Conditioned medium from treated glial cultures is passed through a 30 kDa molecular weight cut off filter, then the concentrated retentate or filtrate is transferred to neurones. As a control, RSL3, erastin or LPS are added to vehicle-treated glial conditioned medium after it is removed from the mixed glial culture. **(b)** Survival of neurones (MTT reduction) relative to concentration of glial conditioned medium. 0 represents glial conditioned medium without LPS treatment; 1 represents neat glial conditioned medium collected from LPS-treated glia; 4 represents 30 kDa MWCO filter retentate diluted into treatment medium to a final concentration equivalent to 4-fold concentration of the neat glial conditioned medium from LPS-treated glia (n=8-11). Dashed lines represent 95% confidence intervals for linear regression. **(c)** Survival of neurones (MTT reduction) after treating with 30 kDa MWCO filtrate of conditioned medium from mixed glial cultures treated with RSL3, erastin or LPS (n=5-6).

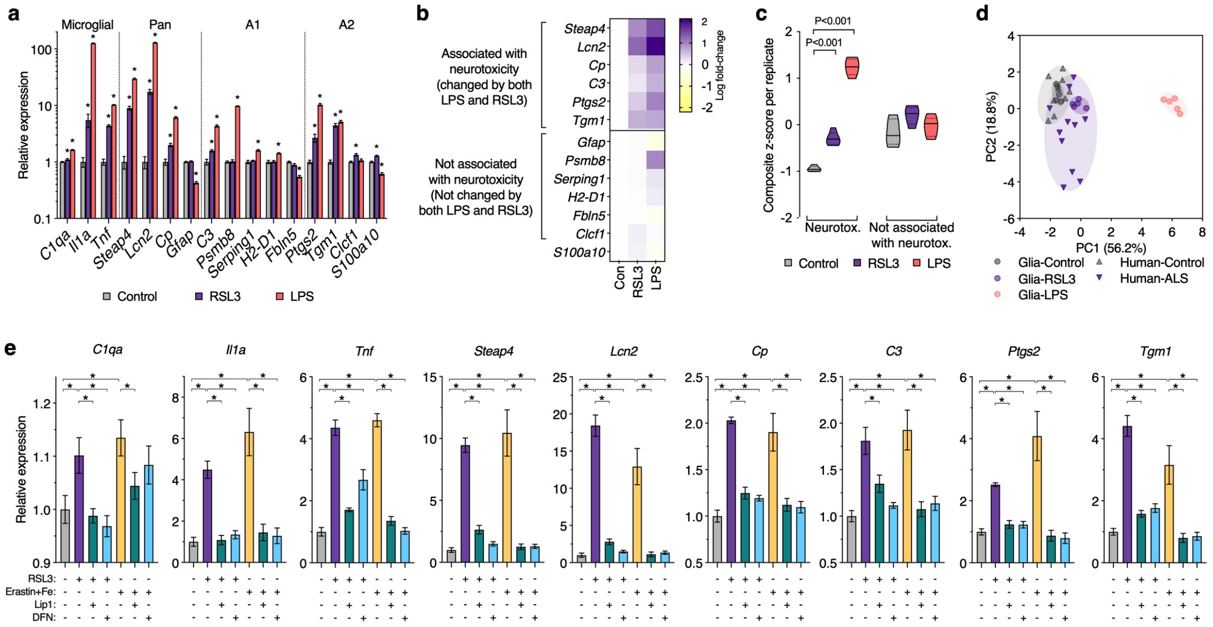


**Supplementary Fig. 6. Gene expression changes in response to ferroptotic stress in glial cells.** **(a)** Expression of genes associated with neurotoxic activation of glia in mixed glial cultures after treatment with RSL3 or LPS (n=5). Selected markers designated for microglial, pan, A1 and A2 activation are highlighted. **(b-c)** Selection of genes used to monitor glial activation in response to ferroptotic stress. Activation genes that changed in response to both stressors and thus associated with neurotoxicity were chosen for subsequent analyses involving glial cultures exposed to LPS or inducers of ferroptotic stress. Violin plots in **c** represent overall transcript signature for features indicated, derived from heatmap data shown in **b** (n=5). **(d)** Principal component analysis of glial activation genes in response to RSL3 or LPS treatment in mixed glial cultures and ALS-affected spinal cord. Symbols represent individual control and ALS cases analysed, or independent mixed glial cultures. Proportion of variance explained by each principal component is denoted on axes. Data are derived from fold expression change for all genes shown in **a** and **Supplementary Fig. 2c**. **(e)** Gene expression changes in mixed glial cultures treated with RSL3- or erastin plus iron (as ferric ammonium citrate) and effects of ferroptosis inhibitors liproxstatin-1 (Lip1) or deferiprone (DFN). These data are presented in heatmap shown in **Figure 3b**. Error margins are S.E.M. (n=3-6). Significant differences indicated by P values (**c**) or asterisks when P<0.05 (**a**,**c**) signifying differences between treatment groups and control (**a**) or as indicated (**e**).

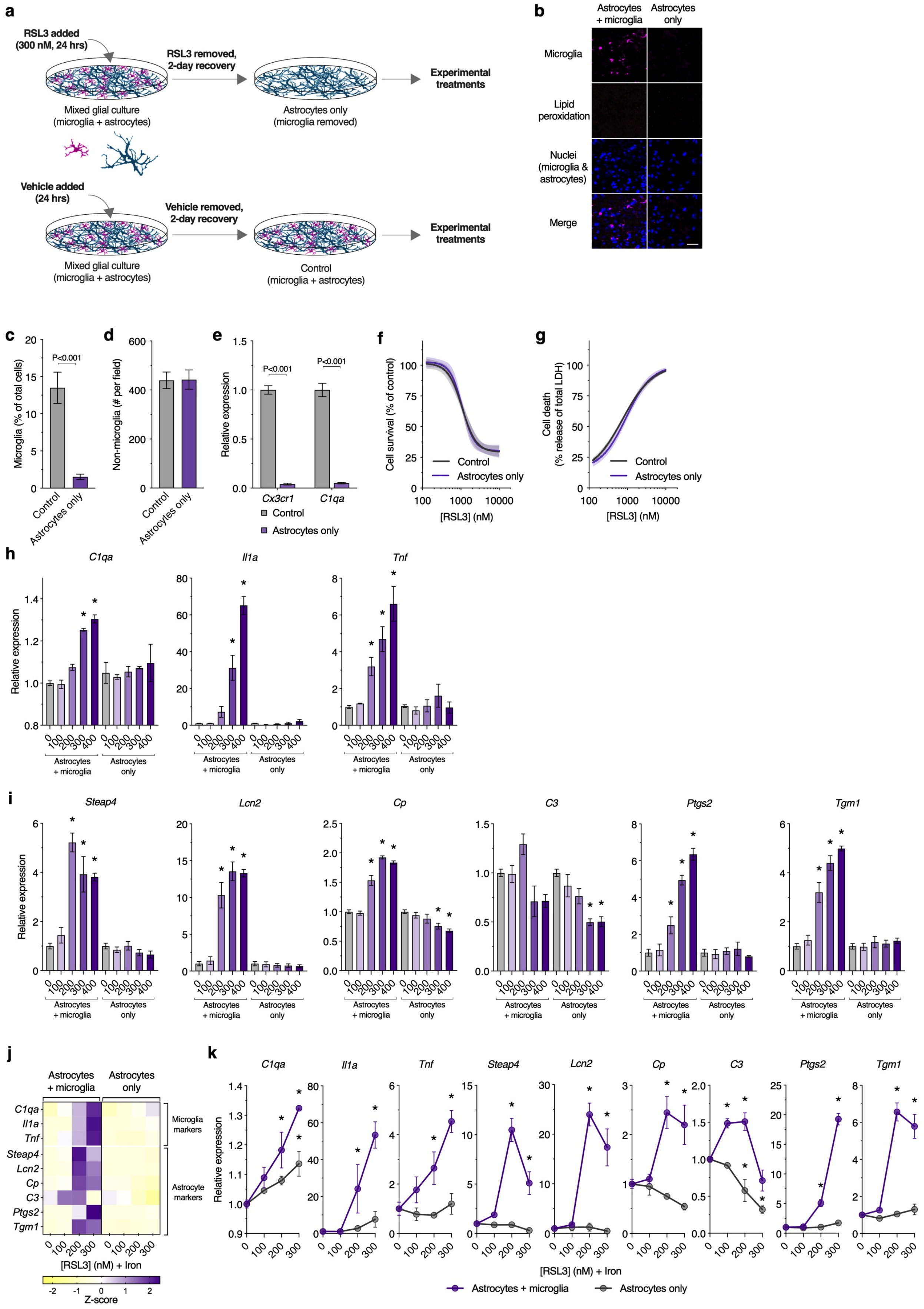


**Supplementary Fig. 7. Isolating astrocytes from mixed glial cultures and gene expression changes in response to treatment with RSL3 or RSL3 plus iron.**

**(a)** Procedure to generate cultures of astrocytes without microglia from mixed glial cultures cultures (containing microglia and astrocytes) by treating with 300 nM RSL3 for 24 hrs. At this RSL3 dose, microglia are killed and astrocytes survive. The RSL3-containing medium is then removed, cultures washed, fresh medium applied, and astrocytes are allowed to recover for 2 days. Parallel treatment of mixed glial cultures with vehicle generates control cultures in which microglia are present. All analyses presented for isolated astrocytes are following RSL3 treatment and the 2-day recovery period. **(b)** Representative fluorescence microscopy images showing microglia and nuclei in control mixed glial cultures and cultures with microglia removed. Lipid peroxidation using C11-BODIPY is non-detectable at the end of the 2-day recovery period. Microglia are depicted as magenta, lipid peroxidation (oxidized:reduced C11-BODIPY) is yellow, and nuclei are blue. Scale bar = 50 µm. **(c-d)** Number of microglia (**c**) or non-microglial cells (**d**) as percentage of total cell population in mixed glial (control) or astrocyte cultures (n=9). **(e)** Expression of the microglial markers *Cx3cr1* and *C1qa* in mixed glial (control) or astrocyte cultures (n=6). **(f,g)** Cell survival (MTT reduction) and cell death (LDH release) in mixed glial (control) or astrocyte cultures in response to RSL3. These data show that the relatively low dose of RSL3 used to remove microglia does not alter subsequent sensitivity of the glial culture to RSL3 after the 2 day recovery period (n=4). **(h,i)** Gene expression changes in mixed glial and astrocyte cultures in response to indicated concentrations of RSL3. These data are presented in heatmap shown in **Figure 3c** (n=3-6). **(j,k)** Expression of genes associated with microglial and astrocyte activation in response to indicated concentrations of RSL3 and transferrin-bound iron in mixed glial or astrocyte cultures (n=3-4). Individual heatmap values in **j** represent mean. Error margins are S.E.M. or 95% confidence intervals for fitted curves (**f**, **g**). P values indicate significant differences between treatment groups as indicated. Asterisks (denoting P<0.05) indicate significant differences between the indicated group and the respective no RSL3 group (**h**, **i**, **k**).

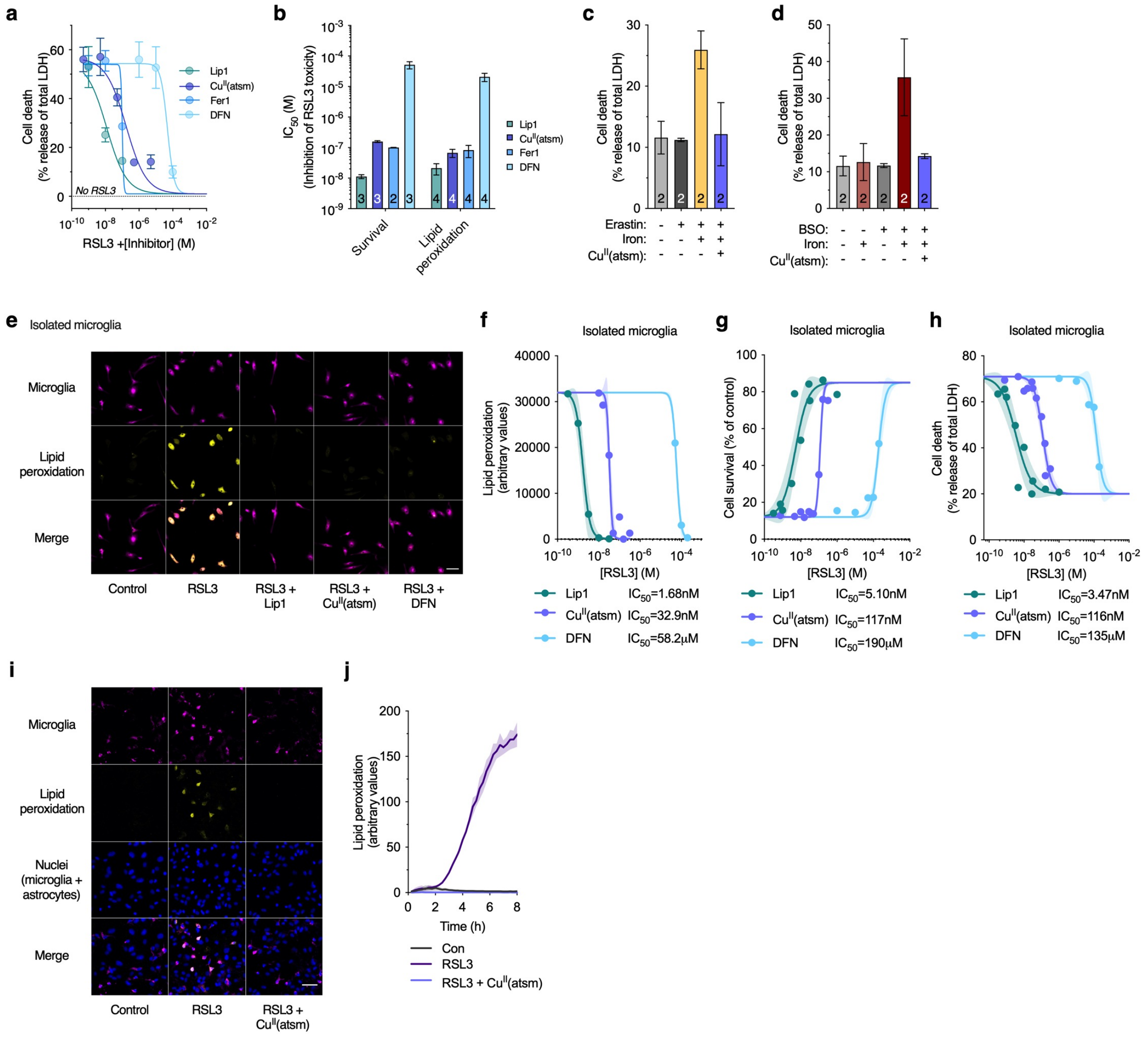


**Supplementary Fig. 8. Protective activity of $Cu^{II}$(atsm) *in vitro*.**

**(a)** $Cu^{II}$(atsm) prevents RSL3-induced cell death (LDH release) in mixed glial cultures, with efficacy similar to the ferroptosis inhibitor ferrostatin-1 (Fer1) (n=3). **(b)** $IC_{50}$ values (derived from data in **Supplementary Fig. 8a** and **Figure 5b**) for liproxstatin-1 (Lip1), $Cu^{II}$(atsm), Fer1 and deferiprone (DFN) against RSL3. **(c,d)** $Cu^{II}$(atsm) prevents cell death (LDH release) in mixed glial cultures treated with ferroptosis inducers erastin plus iron and BSO plus iron (as ferric ammonium citrate). **(e)** Lipid peroxidation (visualised in **Supplementary Video 4**) in microglial cultures treated with RSL3 detected using C11-BODIPY (yellow), is mitigated by Lip1, $Cu^{II}$(atsm) and DFN. **(f-h)** Lipid peroxidation, cell survival (MTT reduction) and cell death (LDH release) in microglial cultures treated with RSL3 and Lip1, $Cu^{II}$(atsm) or DFN. **(i,j)** Microglial lipid peroxidation in mixed glial cultures treated with RSL3 detected using C11-BODIPY (yellow), is mitigated by $Cu^{II}$(atsm). Time dependent changes in microglial lipid peroxidation (j) derived from **Supplementary Video 5** (n=1-3). Error margins are S.E.M. or 95% confidence intervals for fitted lines (**f-h**). Scale bars (**e**, **i**) = 50 μm. Numbers in bars represent number of independent biological replicates.

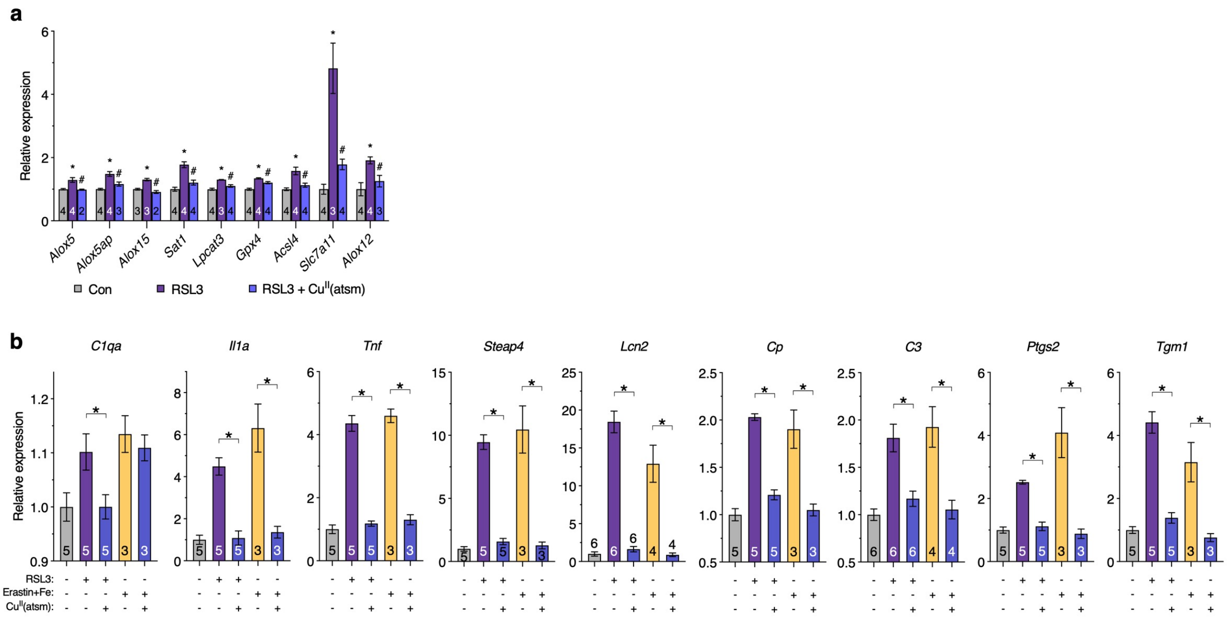


**Supplementary Fig. 9. Gene expression changes in mixed glial cultures treated with inducers of ferroptosis and $Cu^{II}$(atsm).**

**(a)** Relative expression changes for individual genes shown in heatmap in **Figure 5c**. **(b)** Relative expression changes for individual genes shown in heatmap in **Figure 5d**. Asterisks illustrate significant differences (P<0.05) between RSL3 and control (**a**) or indicated treatment groups (**b**). Hash symbols represent significant differences (P<0.05) between RSL3 treated with or without $Cu^{II}$(atsm). Error margins are S.E.M. Numbers in bars represent number of independent biological replicates.

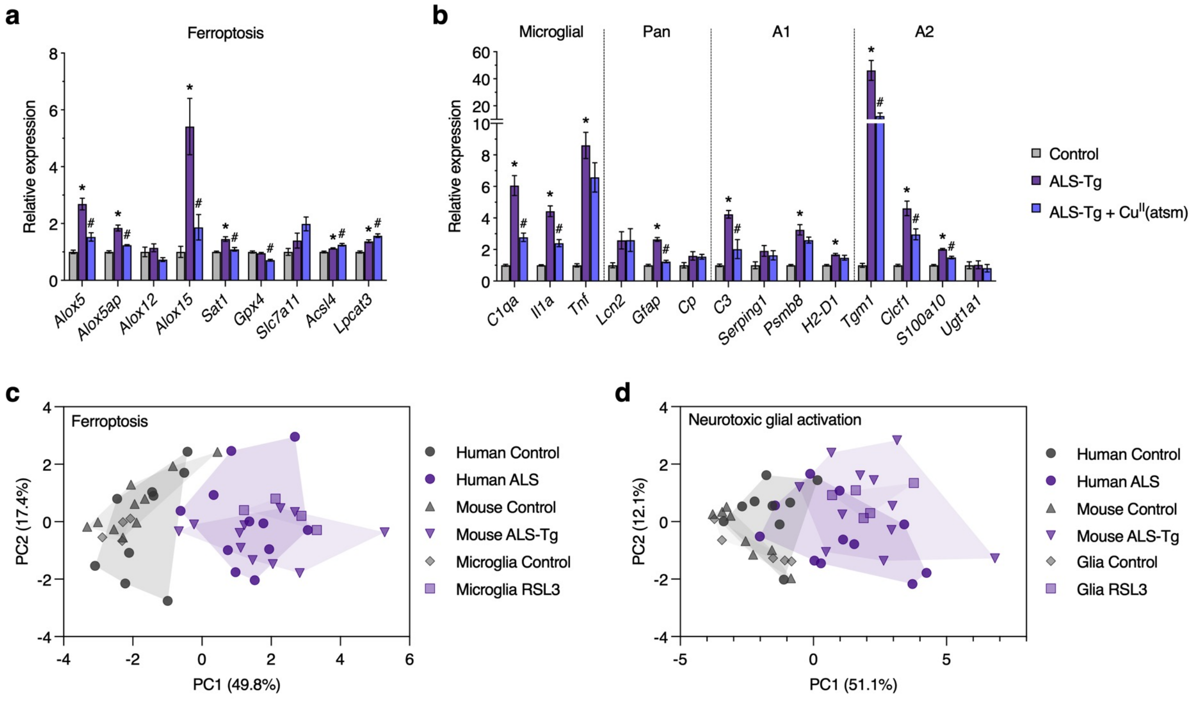


**Supplementary Fig. 10. Gene expression changes in SOD1$^{G37R}$ mice compared to human ALS-affected spinal cord and cultured glia.**

**(a,b)** Relative expression changes for genes associated with (**a**) ferroptosis and (**b**) glial activation, with the latter highlighting selected markers for microglial, pan A1 and A2 activation (n=7-11). These data are presented in heatmaps shown in **Figure 6h** and **6j** respectively. Asterisks illustrate significant differences (P<0.05) between SOD1$^{G37R}$ (ALS-Tg) and respective control. Hash symbols represent significant differences (P<0.05) between SOD1$^{G37R}$ treated with or without Cu$^{II}$(atsm). Error margins are S.E.M. **(c-d)** Principal component analysis of ferroptosis (**c**) and glial activation genes (**d**) in ALS-affected spinal cord, SOD1 $^{G37R}$ mice and glial cultures in response to RSL3 treatment. Symbols represent individual control and ALS cases analysed, individual animals or independent glial cultures. Proportion of variance explained by each principal component is denoted on axes. Data are derived from z-scores of expression changes shown in **Supplementary Fig. 2b**,**c**, **4b**, **6a** and **10a**,**b**.
